# Embryonic Spinocerebellar Ataxia Type 37 AUUUC Repeat RNA Causes Neurodevelopmental Defects in Zebrafish

**DOI:** 10.1101/2025.08.26.672304

**Authors:** Ana F. Castro, Ana S. Figueiredo, Joana R. Loureiro, Maria M. Azevedo, Paula Sampaio, Ana M. Valentim, José Bessa, Isabel Silveira

**Author notes:** Corresponding authors: Isabel Silveira, PhD Genetics of Cognitive Dysfunction Laboratory, i3S - Instituto de Investigação e Inovação em Saúde and IBMC - Institute for Molecular and Cell Biology, University of Porto, Portugal, Rua Alfredo Allen, 208, 4200-135 Porto, Portugal, José Bessa, PhD Vertebrate Development and Regeneration Laboratory, i3S - Instituto de Investigação e Inovação em Saúde and IBMC - Institute for Molecular and Cell Biology, University of Porto, Portugal, Rua Alfredo Allen, 208, 4200-135 Porto, Portugal. Equal contribution (José Bessa, Isabel Silveira).

## Abstract

Onset of many neurodegenerative and neuromuscular diseases usually starts in adulthood; however, recent advances point toward neurodevelopmental changes as drivers of late neurodegeneration. How early neuropathological features occur in these conditions remains unclear, which is critical for timely therapeutic intervention. Here, we provide evidence that neurodevelopmental axonal defects initiate a motor phenotype in a zebrafish model of spinocerebellar ataxia type 37 (SCA37), a degenerative hereditary condition caused by an ATTTC repeat in the *DAB1* gene. We investigated neuronal defects triggered by the embryonic AUUUC repeat RNA and their effects later in life by transiently expressing this RNA in embryos and analyzing innervation and motor function. We found abnormalities in motor neuron axonal outgrowth and muscle innervation. We also discovered disrupted embryonic motor activity and reduced locomotor distance and velocity in late adult zebrafish, demonstrating motor impairment. Moreover, we showed that NOVA2 expression rescues axonal defects, indicating dysfunction of NOVA2-regulated neurodevelopmental processes. Overall, our results establish embryonic expression of the AUUUC repeat RNA as a driver of axonal and synaptic abnormalities, interfering with neuronal circuits and culminating in adult motor dysfunction.

## Introduction

Growing evidence from mouse models of neurodegenerative diseases suggests that abnormal synaptic connectivity during the first days of development sets the stage for neurodegeneration (Capizzi et al., 2022, Edamakanti et al., 2018, Serra et al., 2006, Shabani and Hassan, 2023). This neurodevelopmental dysfunction has for a long time been particularly obvious for spinocerebellar ataxia type 1 (SCA1), where the early developmental expression of the mutant protein consistently leads to abnormal cerebellar connectivity (Edamakanti et al., 2018, Serra et al., 2006). SCAs are autosomal-dominant neurodegenerative diseases characterized by progressive motor incoordination and cerebellar Purkinje cell degeneration (Klockgether et al., 2019). They are frequently caused by an expanded short tandem repeat, or microsatellite sequence, within coding or non-coding gene regions, whose size partly explains the variability in age of disease onset (Loureiro et al., 2022). In a previous study, we have identified an ATTTC repeat inserted in a preexistent polymorphic ATTTT repeat located in an intron of the *DAB1* gene as the cause of SCA37 (Seixas et al., 2017). While the non-pathogenic alleles in the *DAB1* gene may be even larger, ranging from 7 to 400 ATTTT repeat units, they never contain the pathogenic ATTTC repeat insertion (Seixas et al., 2017). *DAB1* is expressed during embryonic development and encodes the reelin adaptor protein, which is known to have an important role in neurodevelopment, including neuronal migration, brain layer organization, and axonal pathfinding (Beffert et al., 2002, D’Arcangelo et al., 1995, Howell et al., 1997, Rice et al., 1998, Sheldon et al., 1997, Trommsdorff et al., 1999). Individuals with SCA37 usually present a phenotype of cerebellar ataxia, including dysarthria and gait and limb incoordination (Corral-Juan et al., 2018, Rosenbohm et al., 2022, Seixas et al., 2017); however, lower motor neuron impairment has also been reported (Rosenbohm et al., 2022), similar to other SCAs (Garcia-Murias et al., 2012, Ikeda et al., 2012, Klockgether et al., 2019). Neuropathological analysis of SCA37 post-mortem cerebellar tissue has shown loss of Purkinje cells, diminished thickness of the molecular cell layer and axonal defects (Corral-Juan et al., 2018). Additionally, a mispositioning of Purkinje cell somas within the molecular and granular cell layers has been observed in brain tissue from elderly SCA37 subjects (Corral-Juan et al., 2018), implying an impairment in proper cerebellar neurodevelopment. This, together with our previous evidence of developmental toxicity of the (ATTTC)_n_ (Seixas et al., 2017), raises the hypothesis that SCA37 may have an early developmental origin similar to what has been found for Huntington’s disease human fetuses (Barnat et al., 2020). Notwithstanding the neuropathological characterization of elderly SCA37 individuals after many decades of disease progression, little is known regarding the embryonic developmental impact of the AUUUC repeat RNA in the disease.

We have shown that the pathogenic *DAB1* (ATTTC)_58_ triggers nuclear RNA aggregation in human cells and causes lethality in zebrafish embryos upon RNA microinjection, indicating that a pathogenic RNA-mediated mechanism contributes to SCA37 (Seixas et al., 2017). Nuclear (AUUUC)_n_ aggregates have been seen in cerebellar Purkinje cells and cortical neurons from brain tissue of affected individuals with a different disease, classified as familial adult myoclonic epilepsy (FAME) (Ishiura et al., 2018). These FAME disorders and SCA37 belong to the rapidly increasing group of currently eight (ATTTC)_n_ insertion conditions (Corbett et al., 2019, Florian et al., 2019, Ishiura et al., 2018, Ishiura and Tsuji, 2020, Loureiro et al., 2022, Silveira and Bennett, 2023, Yeetong et al., 2024, Yeetong et al., 2019), likely sharing an RNA-mediated mechanism. One of the pathogenic processes that seems to explain the cellular and neurological phenotypes associated with the expanded RNA repeats is the abnormal binding and sequestration of RNA-binding proteins (RBPs), which consequently trigger the formation of nuclear RNA aggregates and loss-of-function of the imprisoned RBP in disease (Loureiro et al., 2022, Zhang and Ashizawa, 2022). This mechanism has been demonstrated to play a role in several neurodegenerative conditions caused by non-coding expanded repeats that, as SCA37, result from the expression of RNAs harboring pathogenic repeats (Ishiguro et al., 2017, Swinnen et al., 2018, Todd et al., 2014, White et al., 2010). NOVA2 is an RBP known to have major roles in neurodevelopment, as its dysfunction has been associated with neurodevelopmental disorders (Mattioli et al., 2020, Scala et al., 2022). It is known to control axon guidance and, importantly, control alternative splicing of many neurodevelopmental genes (Ruggiu et al., 2009, Saito et al., 2016, Saito et al., 2019), including the *DAB1* (Corral-Juan et al., 2018, Mattioli et al., 2020, Yano et al., 2010). The very early fetal expression of the AUUUC repeat RNA and its incontestable role in pathogenesis (Seixas et al., 2017) render the investigation of whether and how early neuropathological defects contribute to SCA37 fundamental to advancing knowledge on pathogenesis and timely therapeutic intervention for pentanucleotide repeat and other similar neurodegenerative diseases.

In this study, we established that the embryonic expression of the SCA37 (AUUUC)_58_ RNA in zebrafish leads to developmental axonal and synaptic defects. We showed that the embryonic developmental (AUUUC)_58_ expression disrupts adult zebrafish locomotor function. In addition, we demonstrated that axonal defects are rescued by NOVA2, showing that NOVA2 dysfunction plays a role in SCA37. In summary, we presented developmental motor neuron pathology due to NOVA2 loss-of-function in SCA37, indicating that early therapeutic intervention is key to mitigating this and similar pentanucleotide repeat pathologies.

## Results

### The pathogenic (AUUUC)_58_ RNA induces developmental defects in zebrafish

The SCA37 (AUUUC)_n_ RNA, which is encoded in a *DAB1* intron, is more actively transcribed during embryonic development than in adulthood (Seixas et al., 2017). Therefore, to investigate whether the pathogenic (AUUUC)_n_ RNA affects early vertebrate development, we microinjected 1 to 2-cell stage wild-type zebrafish embryos with the non-pathogenic (AUUUU)_7_ and the pathogenic (AUUUU)_57_(AUUUC)_58_(AUUUU)_>55_ insertion, hereby designated as (AUUUC)_58_, transcribed RNAs, which contain the flanking AluJb monomers as illustrated in **Fig. 1A**, and performed a detailed phenotyping of embryos. We found that the microinjection of the (AUUUC)_58_ RNA significantly increased the lethality and number of morphological developmental defects (77±21%) at 24 hours post-fertilization (hpf), compared with the non-pathogenic (AUUUU)_7_ and vehicle counterparts (33±12% and 25±13%, respectively; p<0.0001; **Fig. S1A**), reproducing our previous findings (Seixas et al., 2017). This shows that the pathogenic (AUUUC)_58_ RNA impairs vertebrate development, as previously demonstrated by our results (Seixas et al., 2017). We also monitored the survival and hatching of the microinjected embryos daily from 0 to 96 hpf. We detected a significantly decreased survival rate (25%) and number of hatched embryos (37%) in the (AUUUC)_58_-injected group, compared with the non-pathogenic (AUUUU)_7_-injected or vehicle group (survival, 55% and 56%, respectively; hatching, 74% and 78%, respectively; p<0.0001 for survival and hatching; **Fig. 1B,C**). This demonstrates that the pathogenic (AUUUC)_58_ RNA decreases the survival of zebrafish embryos, delaying the hatching process. We further examined the morphology of the head and tail in surviving embryos at 24-26 hpf. We established a phenotypical score ranging from 0 to 3 (**Fig. 1D** and **Fig. S1B**), and quantified the number of embryos in each score, finding that the distribution of scores was significantly different among conditions (p<0.0001; **Fig. 1E** and **Fig. S1C**). We found that the microinjection of the pathogenic (AUUUC)_58_ RNA significantly decreased the number of embryos with normal morphology (score 0, 47.22±27.48%), increased the presence of tail and/or head malformations (score 1, 24.63±15.61%; score 2, 26.99±16.28%) and caused development arrest of the zebrafish (score 3, 1.16±1.21%), compared to the non-pathogenic (AUUUU)_7_ RNA (score 0, 89.53±10.40%; score 1, 7.90±8.55%; score 2, 2.39±2.43%; score 3, 0.18±0.46%) or vehicle (score 0, 98.81±1.51%; score 1, 0.38±1.01%; score 2, 0.67±0.94%; score 3, 0.14±0.37%) microinjections (p<0.0001; **Fig. 1E** and **Fig. S1C**). Because pronounced developmental abnormalities have not so far been found in SCA37 family members (Corral-Juan et al., 2018, Rosenbohm et al., 2022, Seixas et al., 2017), and to avoid indirect phenotypes arising from general developmental abnormalities, we selected scored 0 embryos with no visible developmental malformations properly staged as 24 hpf, according to the established morphological criteria of body axis straightening from its early curvature around the yolk (Kimmel et al., 1995), to better characterize the impact of the pathogenic RNA on embryonic development, investigated in the experiments described hereafter.

**Fig. 1.**
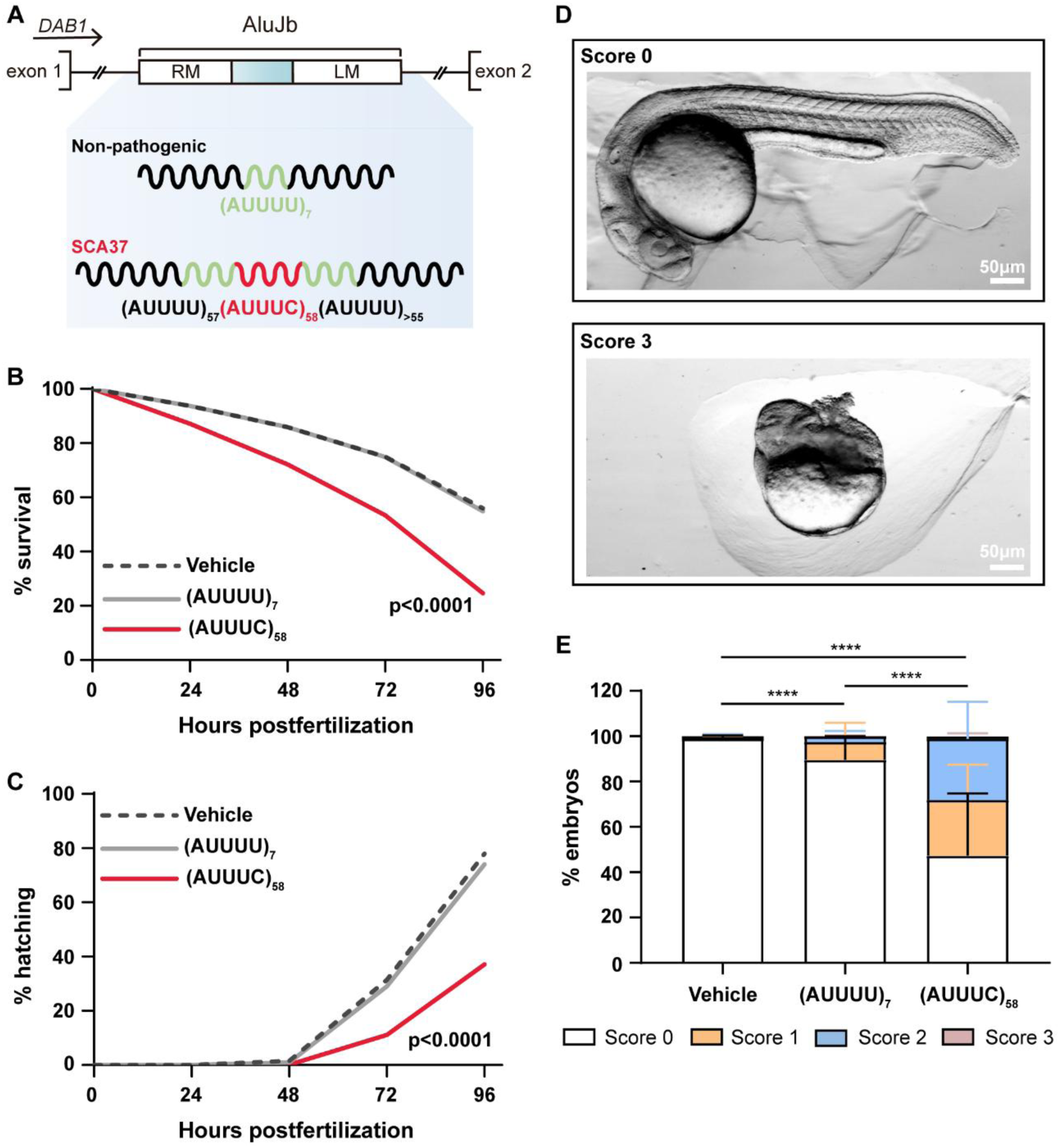
SCA37 (AUUUC)_58_ RNA induces developmental defects. Data presented here were obtained after embryo microinjection of vehicle, (AUUUU)_7_, or (AUUUC)_58_ RNAs, comprising an average of seven independent experiments with at least 100 embryos per experiment. (A) A schematic representation shows the microinjected RNAs, which include the non-pathogenic (AUUUU)_7_ and the pathogenic (AUUUC)_58_ insertion, along with the AluJb flanking regions. (B) Survival rate from 0 to 96 hpf was assessed following microinjection of vehicle, (AUUUU)_7_, or (AUUUC)_58_ RNAs. Statistical significance was determined, revealing notable differences (****p<0.0001, Log Rank test for survival). (C) Hatching rate was assessed from 0 to 96 hpf following the microinjection of vehicle, (AUUUU)_7_, or (AUUUC)_58_ RNAs. The results indicated a significant difference among the conditions (****p<0.0001, Log Rank test for hatching rate). (D) Representative images illustrating the phenotypical scoring scale for embryos are provided, with a score of 0 indicating embryos that are similar to wild-type, while a score of 3 represents the most severe developmental arrest observed. (E) The percentage of embryos categorized by their phenotypical scores at 24 hpf is shown. Scores include score 0 (no morphological defects; white), score 1 (tail defects; orange), score 2 (head and tail defects; blue), and score 3 (developmental arrest; red). The results indicated significant differences across the groups (****p<0.0001, Kruskal-Wallis test followed by Dunn’s post-hoc test for scored embryos distribution). Data are presented as mean ± standard deviation.

### Transient expression of the SCA37 (AUUUC)_58_ RNA in early zebrafish development does not impair zebrin II-positive Purkinje cell differentiation

A consistent phenotype in many SCAs, observed in post-mortem patient tissues, is a marked reduction in the number of cerebellar Purkinje cells (Klockgether et al., 2019). This phenotype could result from an early neurodevelopmental defect leading to impaired Purkinje cell differentiation, in addition to the late Purkinje cell degeneration due to sustained expression of the pathogenic repeat (Shabani and Hassan, 2023). Despite the structural and gene expression differences between human and zebrafish cerebella, the cerebellum in zebrafish is composed of a three-layer structure with similar granular, Purkinje, and molecular cell layers, like in mammals (Bae et al., 2009). During zebrafish cerebellar development, both Purkinje and granule cell neural progenitors start their differentiation at 72 hpf and the positioning of these cells in a three-layer structure is completed at 120 hpf (Kani et al., 2010). Because human RNA repeats are degraded in the first two days after zebrafish embryo microinjection (Todd et al., 2014), we investigated the induction of early and advanced abnormal neuronal phenotypes by the expression of the (AUUUC)_58_ RNA only in this very narrow embryonic time window. To test whether this transient expression is able to impair Purkinje cell progenitor differentiation, we microinjected 1 to 2-cell zebrafish embryos with the pathogenic (AUUUC)_58_ RNA and with the non-pathogenic (AUUUU)_7_ RNA or vehicle, as controls. We selected and raised the microinjected embryos without gross developmental malformations to analyze Purkinje cells at 120 hpf (**Fig. 2A,B**). In this analysis, we quantified the volume occupied by anti-zebrin II, which is known to label all Purkinje cells in zebrafish (Bae et al., 2009, Lannoo et al., 1991a, Lannoo et al., 1991b), as a proxy for estimating the number of Purkinje cells, finding no significant differences among conditions (vehicle, 105456±36670 µm^3^; (AUUUU)_7_, 111705±34446 µm^3^; (AUUUC)_58_, 121519±24115 µm^3^; **Fig. 2C**). Supporting these results, we further assessed the mean intensity of zebrin II fluorescent signal in larvae from microinjected embryos, finding no significant differences among treatments (vehicle, 7219±5961 a.u.; (AUUUU)_7_, 8352±7337 a.u.; (AUUUC)_58_, 8069±6034 a.u.; **Fig. 2D**). These results suggest that the transient expression of the SCA37 (AUUUC)_58_ RNA in early zebrafish development does not impair zebrin II-positive Purkinje cell differentiation.

**Fig. 2.**
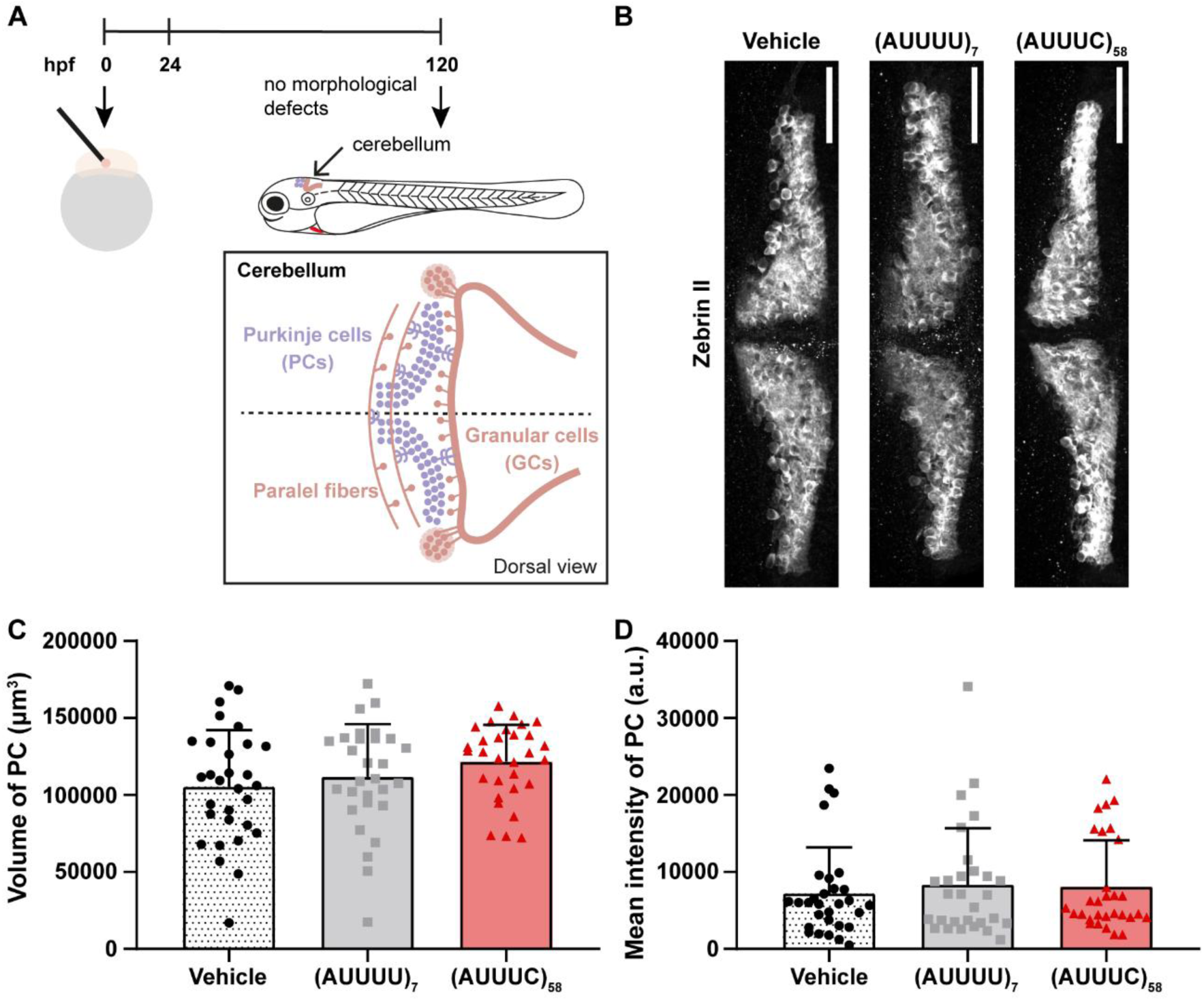
No alterations in zebrin II-positive Purkinje cell differentiation in (AUUUC)_58_ larvae. Data represent three experimental replicates, with a minimum of nine embryos assessed for each condition. (A) The schematic representation outlines the assessment of zebrin II-positive Purkinje cells in zebrafish larvae. One to two-cell zebrafish embryos were microinjected, followed by dead and defective embryos elimination at 24 hpf. Immunofluorescence was conducted using the anti-zebrin II protocol on zebrafish at 120 hpf to analyze zebrin II-positive Purkinje cells. This method allows the visualization and examination of zebrin II-positive Purkinje cell differentiation and organization within the developing nervous system. (B) Representative images depict zebrin II-positive Purkinje cells across each condition. (C) The volume of zebrin II-positive Purkinje cells (µm^3^) was analyzed for each condition. Statistical analysis using a One-way ANOVA indicated no significant differences in the volume of zebrin II-positive Purkinje cells among the conditions. (D) The mean intensity of zebrin II-positive Purkinje cells was evaluated for each condition. Statistical analysis performed using a Kruskal-Wallis test revealed no significant differences in the mean intensity among conditions. Data are presented as mean ± standard deviation. Abbreviations: PC – Purkinje cells; A.U. – arbitrary units.

Because SCA37 patient brain tissue shows cerebellar Purkinje cell loss and a reduction in the thickness of the cerebellar molecular layer (Corral-Juan et al., 2018), where excitatory signals are sent from axons of granule cells (parallel fibers) to Purkinje cell dendrites, we further ruled out the possibility that transient expression of the SCA37 (AUUUC)_58_ RNA during the first hours of development leads to these phenotypes in adulthood. With this in mind, we raised the microinjected embryos and analyzed the cerebellum of adult zebrafish at 19 months of age (**Fig. S2A**). We measured the area of the cerebellar molecular cell layer by staining paraffin longitudinal sections of zebrafish brains with hematoxylin and eosin, and in parallel, we analyzed the number of anti-zebrin II cerebellar Purkinje cells (**Fig. S2B**). We found no significant changes in Purkinje cells arborization by means of cerebellar molecular layer area between (AUUUC)_58_ animals and controls (vehicle, 218761±58368 µm^2^; (AUUUU)_7_, 193656±40226 µm^2^; (AUUUC)_58_, 222280±68435 µm^2^; **Fig. S2C**); however, the total number of zebrin II-positive Purkinje cells was decreased in both the non-pathogenic (AUUUU)_7_ (35±10 cells; p=0.001) and pathogenic (AUUUC)_58_ animals (50±19 cells; p=0.02) when compared with vehicle control animals (80±13 cells; **Fig. S2D**). These findings demonstrate that the transient expression of the SCA37 (AUUUC)_58_ RNA during early zebrafish development has no specific effect on the total number of Purkinje cells. Overall, these results support the idea that the cerebellar phenotypes observed in SCA37 patients, particularly those affecting Purkinje cells, are not caused by early expression of the pathogenic (AUUUC)_58_ RNA alone. Instead, they are rather linked to the degeneration of cerebellar cells due to the sustained expression of the pathogenic RNA repeat.

### The SCA37 (AUUUC)_58_ RNA impairs axonal growth and dorsal-ventral neuromuscular innervation

Since axonal pathology is a hallmark of SCAs (Wilson et al., 2023) and cerebellar axonal degeneration (Corral-Juan et al., 2018) and lower motor neuron impairment (Rosenbohm et al., 2022) have been found in SCA37, we investigated whether the SCA37 (AUUUC)_58_ RNA impairs axonal outgrowth. To test this hypothesis, we analyzed the axons of zebrafish primary motor neurons (PMNs) at 24 hpf (**Fig. 3A** and **Fig. S3A,B**), which exit the spinal cord and extend along each myotome segment towards ventral musculature, using the presynaptic SV2 antibody that recognizes synaptic vesicles (Panzer et al., 2005). We measured the length of PMN axons in 24 hpf zebrafish embryos and observed significantly shorter PMN axons in (AUUUC)_58_-injected embryos (27±8 µm) compared to embryos microinjected with the non-pathogenic (AUUUU)_7_ RNA or vehicle controls (43±12 µm, p=0.0003 and 40±11 µm, p=0.004, respectively; **Fig. 3B**). To rule out any possibility of developmental delay in the (AUUUC)_58_-injected embryos, we compared the axon length of PMNs in wild-type embryos at 19, 22 and 24 hpf (**Fig. S3B,C**). Indeed, we found that the (AUUUC)_58_ RNA microinjection does not impair the initiation of PMN axonal extension but rather slows down its ventral projection (**Fig. S3C,D**). These results show that PMN axonal outgrowth is compromised in the presence of the (AUUUC)_58_ RNA, providing evidence of disturbances in synaptic innervation between spinal cord and muscle tissue. This further suggests that the (AUUUC)_58_ RNA could also impair axonal outgrowth in other neuronal cells. To investigate further whether the (AUUUC)_58_ RNA alters muscle innervation by PMNs and the establishment of the neuromuscular junction (NMJ), we carried out whole-mount immunofluorescence with the presynaptic SV2 antibody and the postsynaptic α-BTX that labels acetylcholine receptors (AchRs) in the muscle (Panzer et al., 2005), at 24 hpf (**Fig. 3C** and **Fig. S4A**). This double staining allowed us to identify NMJs, which are a synaptic connection between the PMNs terminal end and muscle fibers (Panzer et al., 2005). Therefore, we evaluated the co-localization of these two markers along PMN axons and found no clear differences in embryos microinjected with the pathogenic (AUUUC)_58_ RNA compared to those microinjected with the non-pathogenic (AUUUU)_7_ RNA or vehicle (Pearson’s co-localization coefficient 0.3±0.1 in all conditions; **Fig. 3C** and **Fig. S4B**). These results suggest a coupling of pre- and postsynaptic structures between PMN axons and muscle cells in all conditions, indicating that the AchRs distribution in the muscle of (AUUUC)_58_-injected embryos is less extensive in the dorsal-ventral axis, as also observed for PMN axons. To test this hypothesis, we measured the distribution of AchRs in the dorsal and ventral regions of the embryos. Our results showed that the length of AchRs distribution in muscle is shorter in both dorsal and ventral compartments in embryos microinjected with the pathogenic (AUUUC)_58_ RNA (dorsal, 18±8 µm; ventral, 15±8 µm) than in those microinjected with the non-pathogenic (AUUUU)_7_ RNA or vehicle (dorsal, 27±7 µm, p=0.003 and 27±9 µm, p=0.002; ventral, 22±7 µm, p=0.02 and 24±9 µm, p=0.003, respectively; **Fig. 3D**). To investigate potential compensatory mechanisms for the dorsal and ventral loss of AchRs, either through anterior-posterior redistribution or changes in their level, we measured the area and the amount of the AchRs by means of α-BTX signal and its corresponding average intensity, respectively. We observed a decreased area of AchRs (vehicle, 285±70 µm^2^; (AUUUU)_7_, 296±72 µm^2^; (AUUUC)_58_, 203±92 µm^2^; p=0.001, p=0.005; **Fig. 3E**) but no alterations of its fluorescence intensity in pathogenic (AUUUC)_58_ RNA microinjected embryos, compared with controls (vehicle, 335±366 a.u.; (AUUUU)_7_, 385±256 a.u.; (AUUUC)_58_, 207±189 a.u.; **Fig. S4C**). These results demonstrate that the average number of AchRs by area is not different in the dorsal-ventral axis among conditions, showing no changes in terms of AchRs number along the shorter axonal innervation in (AUUUC)_58_ RNA microinjected embryos. Overall, the transient expression of the pathogenic (AUUUC)_58_ RNA impairs axonal growth and dorsal and ventral neuromuscular innervation, suggesting that embryos might develop locomotion defects.

**Fig. 3.**
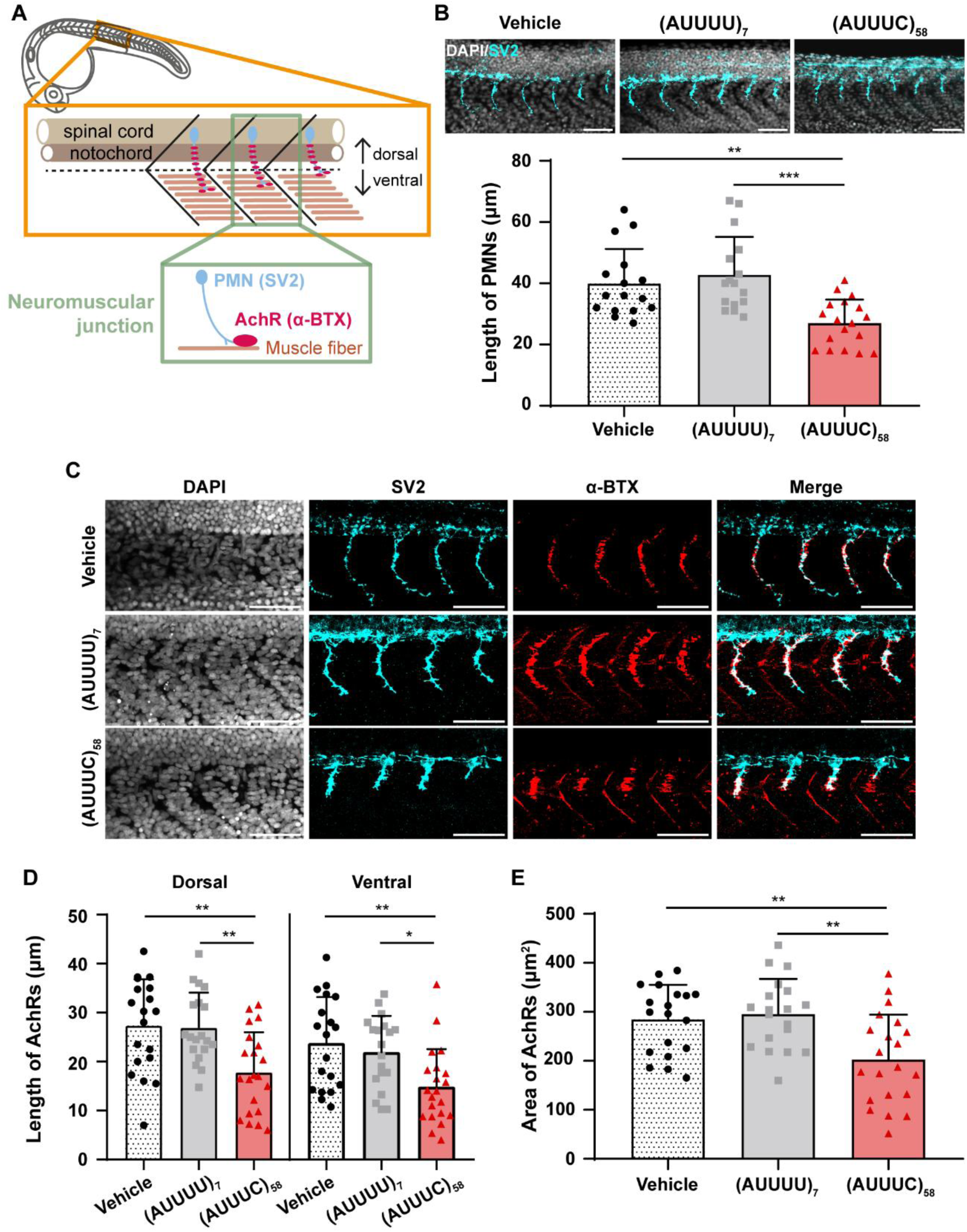
Pathogenic (AUUUC)_58_ RNA induces developmental defects in PMN axons and AchR distribution in muscle. (A) Schematic representation of the analyzed region showing PMN axons innervating muscle cells where AchRs are localized. (B) (top) Representative z-average projection images of PMNs axons in the 6-somites region anterior to the cloaca at 24 hpf (scale bar = 50 µm); (bottom) Quantification of PMN axon lengths in zebrafish embryos microinjected with the vehicle, (AUUUU)_7_ or (AUUUC)_58_ RNAs was performed with the following sample sizes: n=16 embryos for vehicle and (AUUUU)_7_, and n=18 for (AUUUC)_58_ from three experimental replicates. Statistical analysis indicated significant differences (**p<0.01 and ***p<0.001, Kruskal-Wallis test followed by Dunn’s post-hoc test). (C) NMJs are depicted in the 4-somites spanning region anterior to the cloaca at 24 hpf. The images shown are representative z-maximum intensity projections, which highlight the morphology of the NMJs (scale bar = 50 µm). (D) Measurement of AchR signal length along the dorsal-ventral axis of zebrafish muscle was conducted after microinjection of vehicle, (AUUUU)_7_, or (AUUUC)_58_ RNAs. The sample sizes were as follows: n=19 embryos for vehicle and (AUUUU)_7_, and n=21 embryos for (AUUUC)_58_ from three experimental replicates. Statistical analysis revealed significant differences (*p<0.05 and **p<0.01, One-way ANOVA with Bonferroni correction). (E) Quantification of NMJ area (µm^2^) containing AchRs in muscles innervated by PMNs in zebrafish embryos microinjected with the vehicle, (AUUUU)_7_ or (AUUUC)_58_ RNAs (n=19 vehicle and (AUUUU)_7_, n=21 (AUUUC)_58_ from 3 experimental replicates, at 24 hpf; **p<0.01, One-way ANOVA followed by Bonferroni correction). Data are presented as mean ± standard deviation. Abbreviations: PMN – primary motor neuron; AchR – acetylcholine receptor; A.U. – arbitrary units.

### The pathogenic (AUUUC)_58_ RNA disrupts spontaneous early motor activity

Because the microinjection of SCA37 (AUUUC)_58_ RNA leads to defective axonal outgrowth of PMNs and muscle innervation, we inferred that other important neuronal defects might also be induced, leading to motor dysfunction. To test this hypothesis, we tested locomotor activity induced by a touch stimulus and spontaneous motor activity. For the former, we performed the touch-evoked escape swimming response test in previously microinjected larvae with the (AUUUC)_58_ RNA and controls at 72 hpf (**Fig. S5A**). We observed no significant differences in the percentage of larvae responding to touch stimuli among groups (60% for vehicle, 57% for (AUUUU)_7_ and 59% for (AUUUC)_58_; **Fig. S5B**). In larvae that responded to the touch stimulus, we analyzed the time between the stimulus and the larvae response (reaction time) and the duration of the escape response (swimming duration response) for each larva. We found no differences in the reaction time (vehicle, 0.2±0.1 s; (AUUUU)_7_, 0.3±0.2 s; (AUUUC)_58_ 0.2±0.1 s; **Fig. S5C**) or swimming duration response (vehicle, 2.3±1.8 s; (AUUUU)_7_, 2.1±1.8 s; (AUUUC)_58_ 2.0±1.7 s; **Fig. S5D**) of larvae microinjected with the (AUUUC)_58_ RNA when compared with controls. These results indicate that the neuronal alterations at 24 hpf are insufficient to impair the response of larvae to a touch stimulus later at 72 hpf. Next, we tested whether the early transient presence of the pathogenic (AUUUC)_58_ RNA could affect spontaneous motor activity. Specifically, we assessed the number of spontaneous tail coils at 24 hpf and the free-swimming behavior of larvae at 144 hpf. Thus, we measured the number of spontaneous tail coils over a 60-second period (**Fig. 4A**). Spontaneous tail coiling represents the first involuntary movements in embryos and serves as a measure of synaptic communication between spinal cord and muscle cells (Drapeau et al., 2002). We observed a significant increase in tail coiling movements in embryos microinjected with the (AUUUC)_58_ RNA (5±6 coils) compared to those microinjected with the non-pathogenic (AUUUU)_7_ RNA or vehicle controls (3±2 coils, p=0.043 and 3±2 coils, p=0.003, respectively; **Fig. 4B** and **Movies 1-3**), supporting the hypothesis that the (AUUUC)_58_ RNA alters spinal cord-muscle innervation, leading to an aberrant increase in tail movements. Apart from these very early spontaneous movements, we further analyzed the free-swimming behavior of the larvae at 144 hpf previously microinjected with the pathogenic (AUUUC)_58_ RNA, the non-pathogenic (AUUUU)_7_ RNA, or the vehicle. We analyzed the percentage of immobile larvae during the 10-minute test in each group, finding no significant differences among them (37% for vehicle, 39% for (AUUUU)_7_ and 27% for (AUUUC)_58_; **Fig. 4C**). For motile larvae, we measured the distance swum by each larva during the test and found that it tended to be shorter in larvae microinjected with the pathogenic (AUUUC)_58_ RNA compared to the control conditions (vehicle, 50±55 cm; (AUUUU)_7_, 37±50 cm; (AUUUC)_58_, 27±36 cm) (**Fig. 4D**). Concomitantly, we observed that for motile larvae, the average time spent immobile tended to be slightly higher in larvae microinjected with the pathogenic (AUUUC)_58_ RNA, compared to controls (vehicle, 257±222 s; (AUUUU)_7_, 333±227 s; (AUUUC)_58_, 356±202 s; **Fig. 4E**). Considering an immobile episode a 2-second interval in which no locomotor activity was detected, we found that larvae microinjected with the pathogenic (AUUUC)_58_ RNA exhibited an increased number of immobility episodes (33±30 episodes) compared to those microinjected with the vehicle control (14±16 episodes, p=0.02; **Fig. 4F**). Overall, these results indicate that the transient expression of SCA37 (AUUUC)_58_ RNA in early zebrafish embryonic development impairs spontaneous locomotor activity and excitability. Our results showed that SCA37 (AUUUC)_58_ RNA disrupts PMN axonal outgrowth and causes alterations in neuromuscular innervation. Although these changes are unlikely to directly cause the observed behavior phenotypes in spontaneous coiling and free-swimming tests (e.g. PMNs innervate fast muscle fibers that are not involved in zebrafish coiling activity (Naganawa and Hirata, 2011)), similar defects in other neuronal populations may indeed underlie the observed motor alterations.

**Fig. 4.**
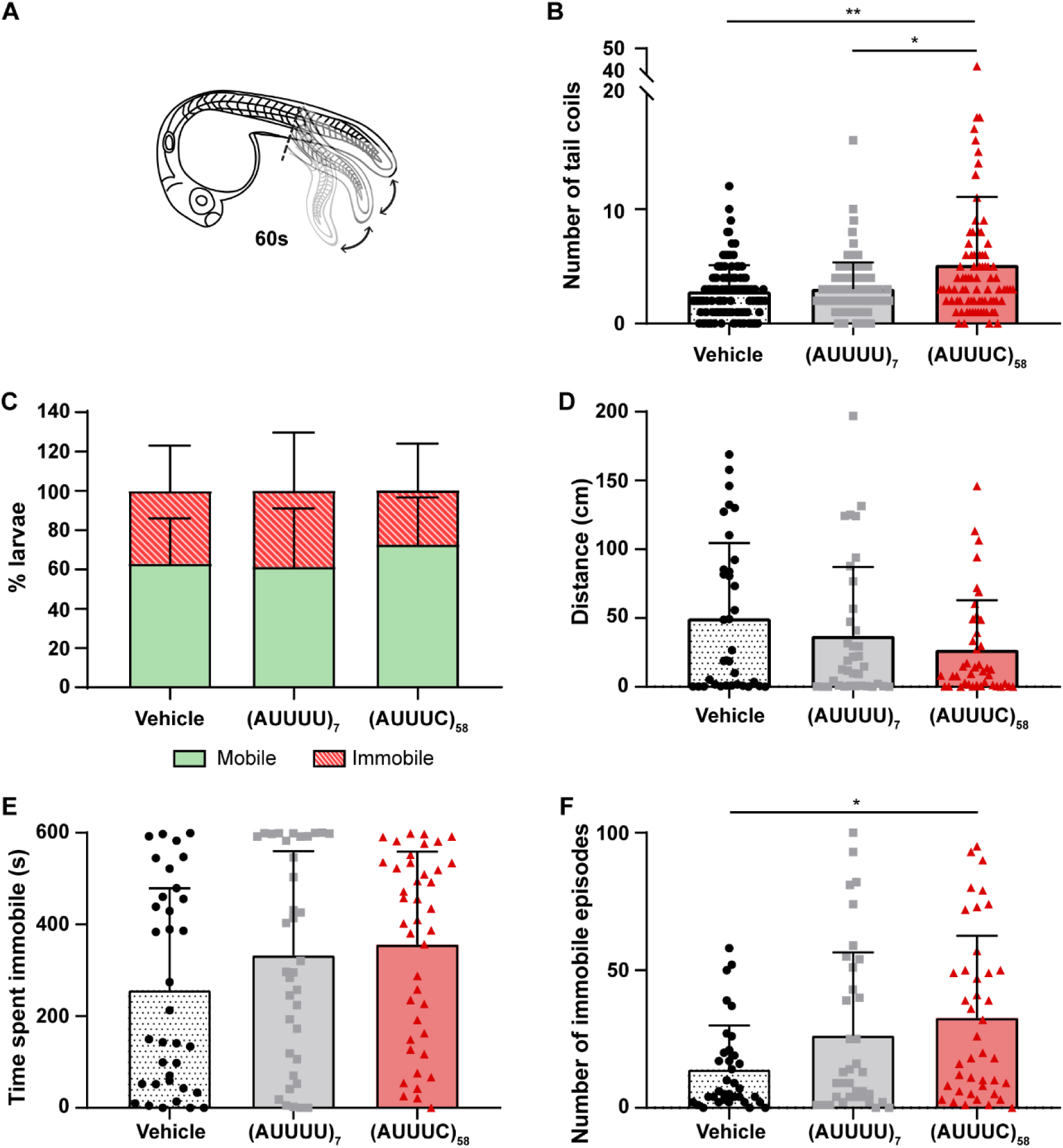
(AUUUC)_58_-injected animals exhibit hyperactivity and frequent episodes of impaired motor function during development. (A) Schematic representation illustrating involuntary tail movements in 24 hpf zebrafish embryos. (B) Quantification of spontaneous tail coil movements over a 60-second period in zebrafish embryos microinjected with vehicle, (AUUUU)_7_ or (AUUUC)_58_ RNAs (n=112 embryos in vehicle and (AUUUU)_7_ groups, n=82 in (AUUUC)_58_ group from seven experimental replicates, at 24 hpf; *p<0.05 and **p<0.01, Kruskal-Wallis test followed by Dunn’s post-hoc test). (C) Percentage of larvae that remained mobile and immobile during a 10-minute spontaneous locomotor test (n=55 vehicle larvae, n=58 (AUUUU)_7_ larvae, n=57 (AUUUC)_58_ larvae from three experimental replicates; ꭕ2 test for mobility or immobility). (D) Total swimming distance was assessed in the spontaneous locomotor test with the following sample sizes: vehicle n=35 larvae, (AUUUU)_7_ n=36 larvae, and (AUUUC)_58_ n=41 larvae across three experimental replicates. Statistical analysis was performed using the Kruskal-Wallis test and no significant differences were found. (E) The time spent immobile by larvae was measured with the following sample sizes: vehicle n=35 larvae, (AUUUU)_7_ n=36 larvae, and (AUUUC)_58_ n=41 larvae from three experimental replicates. Statistical analysis was conducted using the Kruskal-Wallis test, and no significant differences were observed. (F) The number of immobile episodes (lasting more than 2 seconds) in larvae was quantified with the following sample sizes: vehicle n=35 larvae, (AUUUU)_7_ n=36 larvae, and (AUUUC)_58_ n=41 larvae from three experimental replicates. Statistical analysis revealed significant differences between (AUUUC)_58_ and vehicle groups (*p<0.05, Kruskal-Wallis test followed by Dunn’s post-hoc test). Data are presented as mean ± standard deviation.

### Transient expression of the SCA37 (AUUUC)_58_ RNA in early zebrafish development alters locomotor profile in adulthood

Given the alterations found during zebrafish development, we asked whether the transient expression of the SCA37 (AUUUC)_58_ RNA has a long-term impact on zebrafish locomotor function. Therefore, we performed the novel diving tank test at 3 months in animals previously microinjected in the 1 to 2-cell stage with the pathogenic (AUUUC)_58_ RNA, non-pathogenic (AUUUU)_7_ RNA, or vehicle. This test consists of placing the zebrafish into a new tank and recording its movements for 6 minutes during the exploratory phase, allowing the analysis of animal locomotor activity in a new environment. The 3-month-old animals did not show striking differences among conditions in the percentage of time spent immobile (vehicle, 6±3%; (AUUUU)_7_, 8±4%; (AUUUC)_58_, 5±2%; **Fig. S6A**) or the distance swum (vehicle, 23±6 m; (AUUUU)_7_, 23±8 m; (AUUUC)_58_, 28±4 m; **Fig. S6B**). However, the pathogenic (AUUUC)_58_ RNA microinjected animals exhibited a mild increase in displacement velocity (8±1 cm/s) compared to vehicle control animals (7±2 cm/s, p=0.045; **Fig. S6C**). Additionally, no differences in body length and weight of the animals were found among experimental groups (vehicle, 1.95±0.25 cm and 126±94 mg; (AUUUU)_7_, 1.98±0.34 cm and 129±64 mg; (AUUUC)_58_, 2.12±0.23 cm and 166±87 mg; **Fig. S7A,B**). Given that SCA37 is a late-onset disease (Seixas et al., 2017), we performed the novel diving tank test on 17-month-old zebrafish to investigate locomotor dysfunction in late adults. While no significant differences were observed in the percentage of time spent immobile among conditions (vehicle, 6±4%; (AUUUU)_7_, 7±6%; (AUUUC)_58_, 9±10%; **Fig. 5A**), zebrafish microinjected with pathogenic (AUUUC)_58_ RNA exhibited a significant shorter swimming distance (28±9 m) and reduced speed (9±2 cm/s) compared to those microinjected with non-pathogenic (AUUUU)_7_ RNA (39±11 m, p=0.005 and 12±3 cm/s, p=0.0003, respectively; **Fig. 5B,C**), demonstrating a locomotor dysfunction in these animals. Furthermore, we observed no differences in the length and weight of the animals among conditions (vehicle, 2.86±0.19 cm and 585±184 mg; (AUUUU)_7_, 2.95±0.16 cm and 533±113 mg; (AUUUC)_58_, 2.89±0.22 cm and 564±206 mg; **Fig. S7C,D**). When comparing the locomotor performance of 3- and 17-month-old adult zebrafish, no significant differences were observed in the percentage of time spent immobile (vehicle, t_3mth_ = 6±3% and t_17mth_ = 6±4%; (AUUUU)_7_, t_3mth_ = 8±4% and t_17mth_ = 7±6%; (AUUUC)_58_, t_3mth_ = 5±2% and t_17mth_ = 9±10%; **Fig. 6A**); however, the 17-month-old control animals swam farther (vehicle, d_3mth_ = 23±6 m and d_17mth_ = 31±6 m, p=0.003; (AUUUU)_7_, d_3mth_ = 23±8 m and d_17mth_ = 39±11 m, p<0.0001; **Fig. 6B**) and faster (vehicle, v_3mth_ = 7±2 cm/s and v_17mth_ = 9±1 cm/s, p=0.001; (AUUUU)_7_, v_3mth_ = 7±2 cm/s and v_17mth_ = 12±3 cm/s, p<0.0001; **Fig. 6C**), compared to 3-month-old control animals. This increase in performance is expected, as animals typically grow (length and weight) between 3 and 17 months of age (see **Fig. S7**). Although zebrafish microinjected with pathogenic (AUUUC)_58_ RNA exhibited an increase in size as control conditions (**Fig. S6**), they did not show the corresponding increase in locomotor performance (d_3mth_ = 28±4 m and d_17mth_ = 28±9 m; v_3mth_ = 8±1 cm/s and v_17mth_ = 8±2 cm/s; **Fig. 6B,C**). Thus, there was a significant interaction between the factors condition and age (p=0.0008 for distance and p=0.0004 for speed), which shows that these animals experience locomotor impairment that becomes evident in late adulthood.

**Fig. 5.**
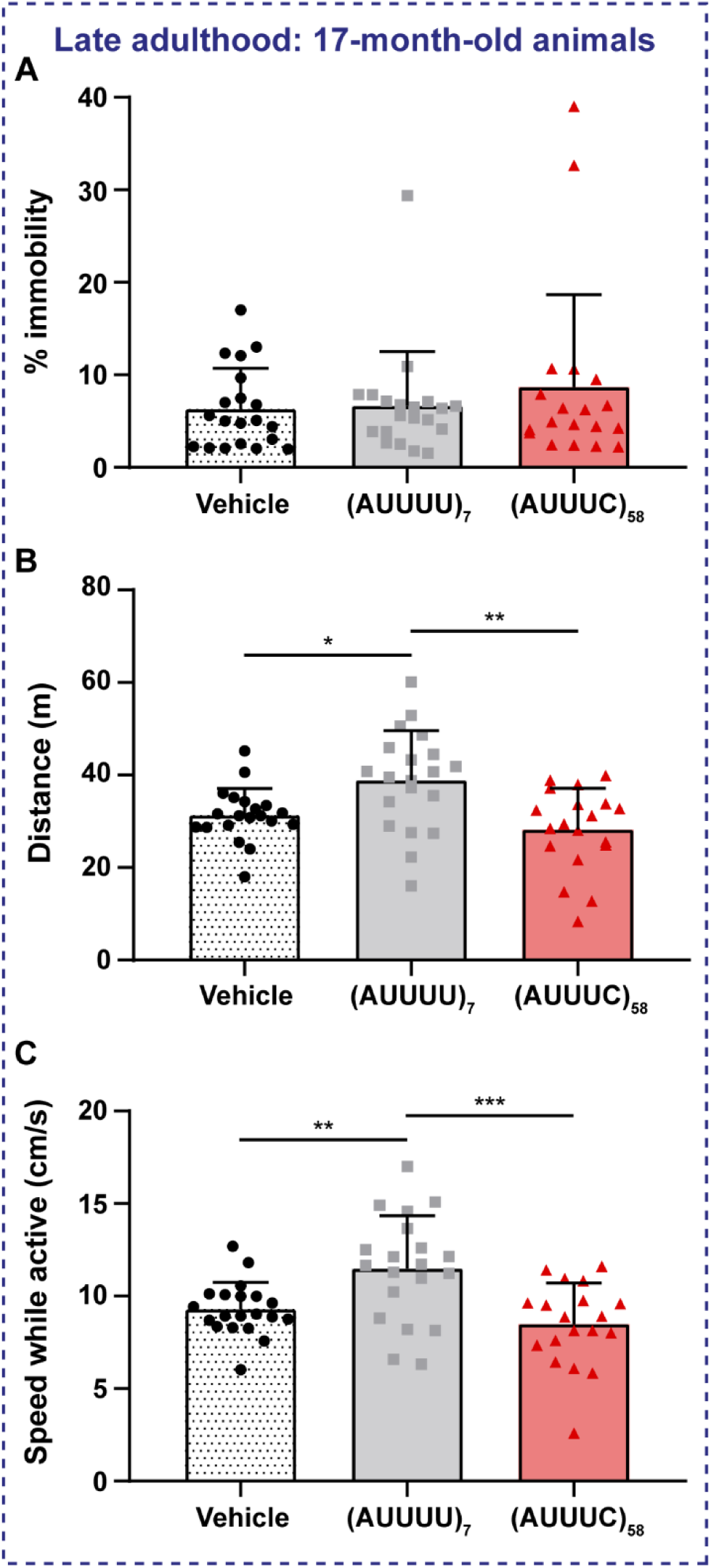
Pathogenic (AUUUC)_58_ causes motor impairment in late-adult zebrafish. The novel diving tank test was conducted on microinjected animals at 17 months of age, n=20 for the vehicle and (AUUUU)_7_ conditions, and n=19 for the (AUUUC)_58_ condition. (A) Analysis of the percentage of immobile animals was conducted using the Kruskal-Wallis test, yielding no significant results. (B) Quantification of the total swimming distance was conducted, yielding results that were statistically significant (*p<0.05 and **p<0.01, One-way ANOVA following Welch correction and Dunnett’s T3 post-hoc test). (C) Quantification of the average speed while animals are swimming (**p<0.01 and ***p<0.001, One-way ANOVA followed by Bonferroni correction). Data are presented as mean ± standard deviation.

**Fig. 6.**
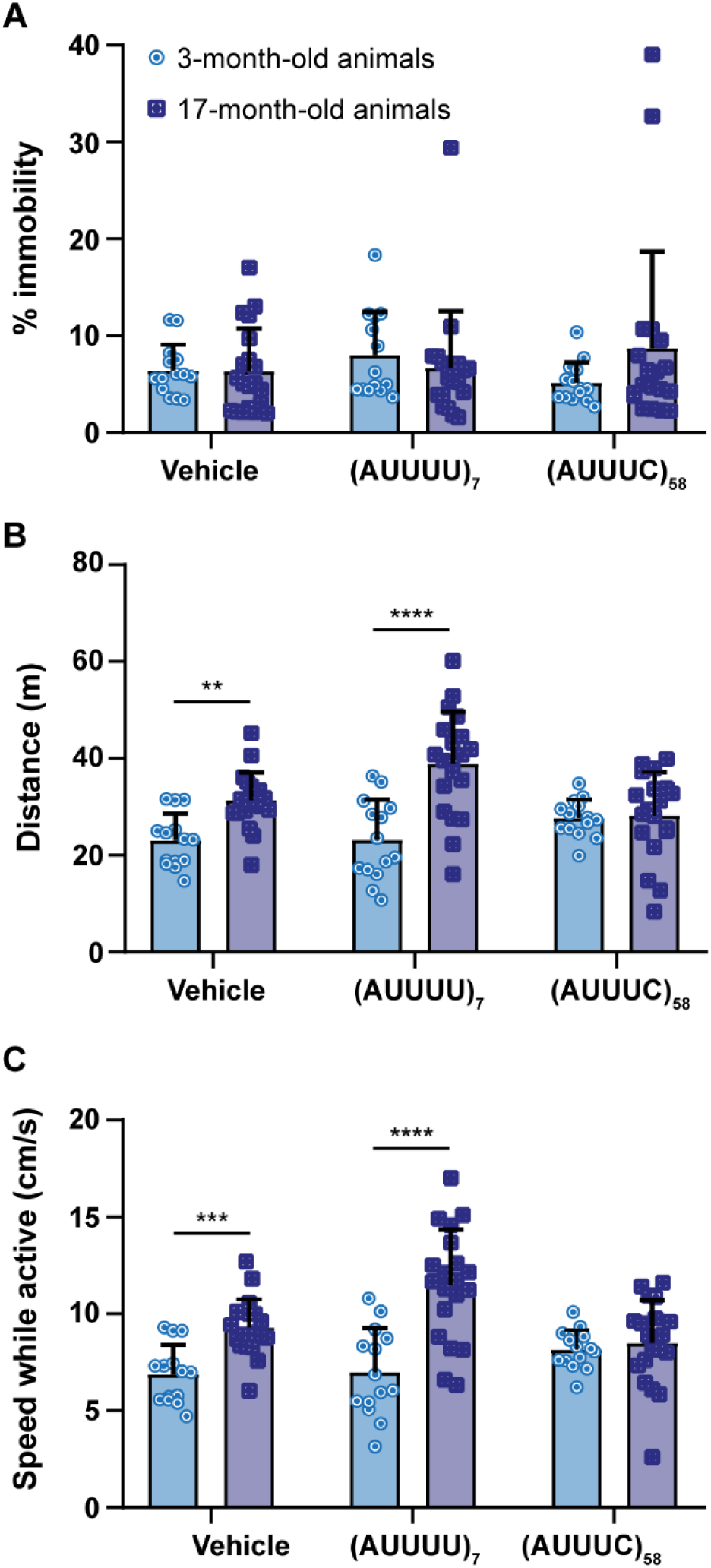
Pathogenic (AUUUC)_58_ RNA negatively affects motor performance from 3 to 17 months. The novel diving tank test was performed at 3 months with n=14 animals per condition and at 17 months with n=20 animals for the vehicle and (AUUUU)_7_ conditions, and n=19 animals for the (AUUUC)_58_ condition. (A) Analysis of the percentage of immobile animals (Mann-Whitney U test). (B) Quantification of the total distance swum by the animals (**p<0.01 and ****p<0.0001, Two-way ANOVA with condition and age as factors for distance). (C) Analysis of the average swimming speed while the animals are active (***p<0.001 and ****p<0.0001, Two-way ANOVA with condition and age as factors for speed). Data are shown as mean ± standard deviation.

### NOVA2 rescues defective axonal outgrowth in zebrafish with the pathogenic (AUUUC)_58_ RNA

One of the mechanisms associated with the pathogenic (AUUUC)_58_ RNA in SCA37 is expected to involve the sequestration of RBPs, leading to their functional impairment. Therefore, we performed an unbiased *in silico* analysis to identify RBPs with high binding affinity to the pathogenic (AUUUC)_58_ RNA, using the RBPmap webserver (Paz et al., 2014). We found 109 putative RBPs predicted to bind to the non-pathogenic (AUUUU)_170_ and pathogenic (AUUUC)_58_ RNAs (p<0.05; **Table S1**). Next, we determined the maximum z-score for each RBP hit, detecting NOVA1 as the unique RBP in this database that binds with more affinity to the pathogenic (AUUUC)_58_ RNA than to the non-pathogenic RNA (**Fig. 7A**). We also quantified the number of binding sites for NOVA1 in both non-pathogenic and SCA37 (AUUUC)_58_ RNAs. We found 61 binding sites in (AUUUC)_58_ RNA, against the 3 binding sites in (AUUUU)_170_ RNA, which has the same total repeat length as the pathogenic RNA with the flanking AUUUU repeats (**Fig. 7B**). NOVA2 RBP, which is not present in the RBPmap database like NOVA1, is a neuron-specific splicing factor that binds to the same YCAY binding motif (Saito et al., 2016) present in the AUUUC repeat itself, as NOVA1. Moreover, NOVA2 plays a role in axonal pathfinding (Saito et al., 2016) and is expressed in the same neuronal cells as DAB1 (Yano et al., 2010). Therefore, we investigated whether NOVA2 loss-of-function might be implicated in the PMN axonal defects detected in zebrafish embryos microinjected with the pathogenic (AUUUC)_58_ RNA. We co-microinjected the pathogenic (AUUUC)_58_ RNA and the human *NOVA2* mRNA into zebrafish embryos and then analyzed PMN axon length at 24 hpf using whole-mount immunofluorescence with anti-SV2 (**Fig. 7C**). We confirmed that *NOVA2* mRNA microinjection alone does not impact the length of PMN axons in zebrafish when compared to vehicle control embryos (vehicle, 43±7 µm; NOVA2, 39±9 µm; **Fig. 7D**), after its translation into NOVA2 protein assessed by whole-mount immunofluorescence with anti-histidine antibody targeting the NOVA2 histidine tail (**Fig. 7E**). As previously observed, we also confirmed that the pathogenic (AUUUC)_58_ RNA microinjection leads to axonal shortening of PMNs (31±8 µm) when compared to control conditions (vehicle, 43±7 µm; (AUUUU)_7_, 41±7 µm; p<0.0001 for both controls; **Fig. 7D**). Importantly, we found that NOVA2 RBP suppresses the axonal shortening of PMNs triggered by the pathogenic (AUUUC)_58_ RNA, rescuing the length of zebrafish PMN axons, when co-microinjected with the (AUUUC)_58_ RNA ((AUUUC)_58_, 31±8 µm; (AUUUC)_58_+NOVA2, 37±8 µm, p=0.01; **Fig. 7C,D**). Although the embryos co-microinjected with the (AUUUC)_58_ RNA and *NOVA2* mRNA exhibited shorter PMN axons ((AUUUC)_58_+NOVA2, 37±8 µm) compared to vehicle control embryos (vehicle, 43±7 µm), their axonal lengths were not significantly different from those observed in the other non-pathogenic conditions (NOVA2, 39±9 µm; (AUUUU)_7_, 41±7 µm; **Fig. 7D**). These results indicate that the SCA37 pathogenic mechanism triggers a partial NOVA2 loss-of-function.

**Fig. 7.**
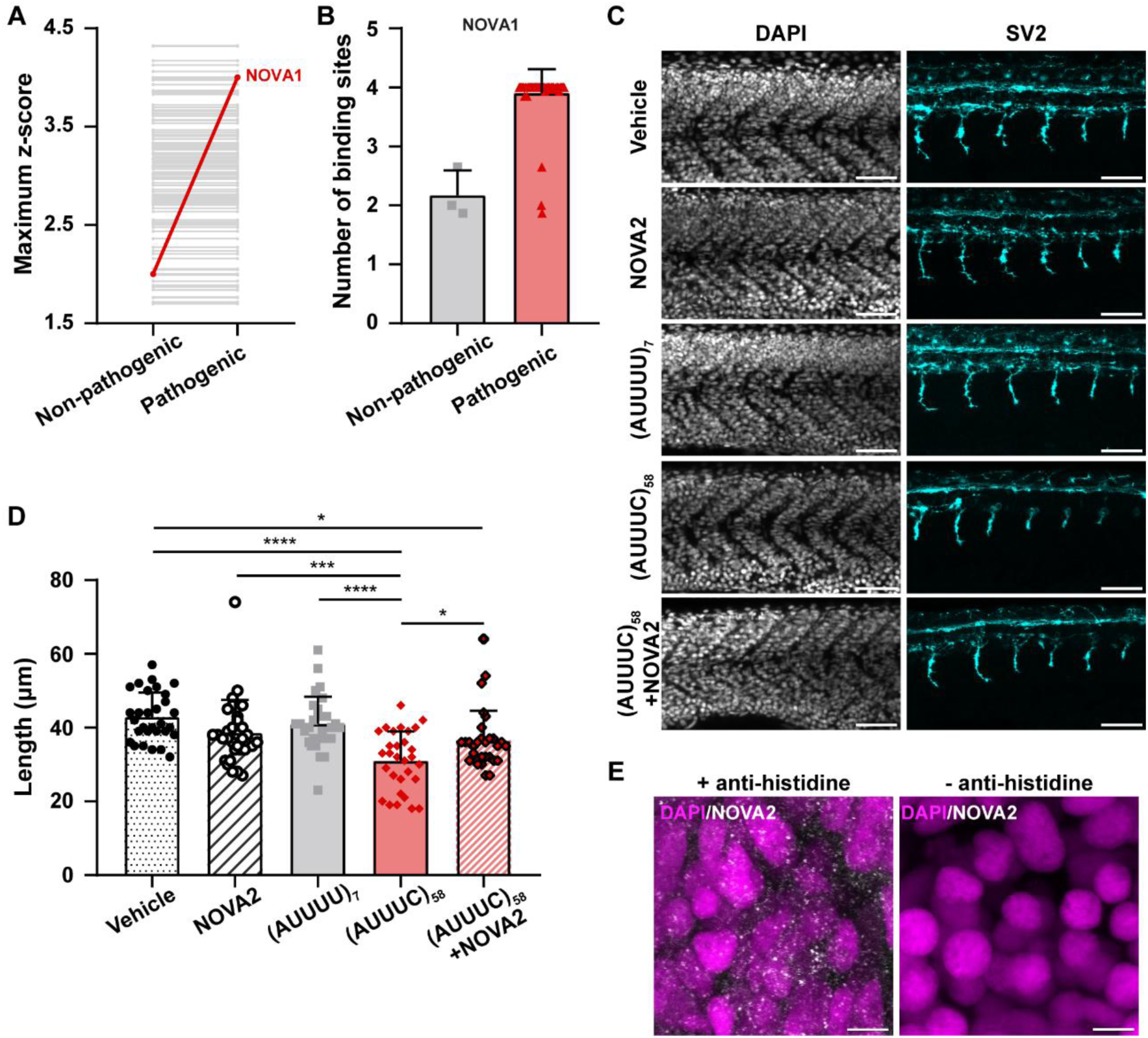
NOVA2 mitigates axonal defects caused by pathogenic (AUUUC)_58_ RNA in PMNs. (A) Maximum z-scores of the putative RBPs that bind to the non-pathogenic (ATTTT)_170_ and pathogenic (AUUUC)_58_ RNAs, assessed by the RBPmap tool (p<0.05). (B) Number of binding sites for NOVA1 in the non-pathogenic (ATTTT)_170_ and SCA37 (AUUUC)_58_ RNAs. Data are shown as mean ± standard deviation. (C) PMN axons located in the 6-somite region anterior to the cloaca at 24 hpf are displayed, featuring representative z-average intensity projection images (scale bar = 50 µm). (D) Length of PMN axons in zebrafish embryos microinjected with either the vehicle, NOVA2, (AUUUU)_7_, (AUUUC)_58_, or a combination of (AUUUC)_58_ and *NOVA2* mRNA (n=30 embryos per condition across 3experimental replicates). Statistical analysis revealed significant differences (*p<0.05, ***p0.001 and ****p<0.0001) using a Two-way ANOVA with conditions and replicates as factors, followed by a Bonferroni post-hoc test for axon length in each condition, after applying cube root data transformation. Raw data are represented as mean ± standard deviation. (E) Observation of nuclear and cytoplasmic distribution of the human NOVA2 protein in zebrafish embryos microinjected with *NOVA2* mRNA in the muscle region anterior to the cloaca (left). The images presented are representative z-maximum intensity projections with (left) and without (right) anti-histidine primary antibody (scale bar = 5 µm).

## Discussion

In this study, we demonstrated that the pathogenic (AUUUC)_58_ RNA embryonically causes axon outgrowth defects in presynaptic PMNs and aberrant postsynaptic localization of AchRs clusters in zebrafish. In our experimental setting, we transiently expressed the SCA37 (AUUUC)_58_ RNA in early zebrafish embryos and demonstrated that the developmental defects caused by this RNA contribute to impaired motor function in adult animals. Furthermore, we demonstrated that co-microinjection of the pathogenic (AUUUC)_58_ RNA with human NOVA2 rescues axon outgrowth defects caused by the (AUUUC)_58_. Thus, we propose that (1) SCA37 pathogenesis may be partially driven by significant neurodevelopment defects caused by the early expression of the pathogenic (AUUUC)_58_ RNA, (2) these neurodevelopmental defects result in locomotor impairment in late adulthood, and (3) a key mechanism underlying these neurodevelopmental defects is the partial loss-of-function of NOVA2 due to its sequestration by the pathogenic (AUUUC)_58_ RNA, which contains an abnormally high number of predicted binding sites for this RBP (**Fig. 8**).

**Fig. 8.**
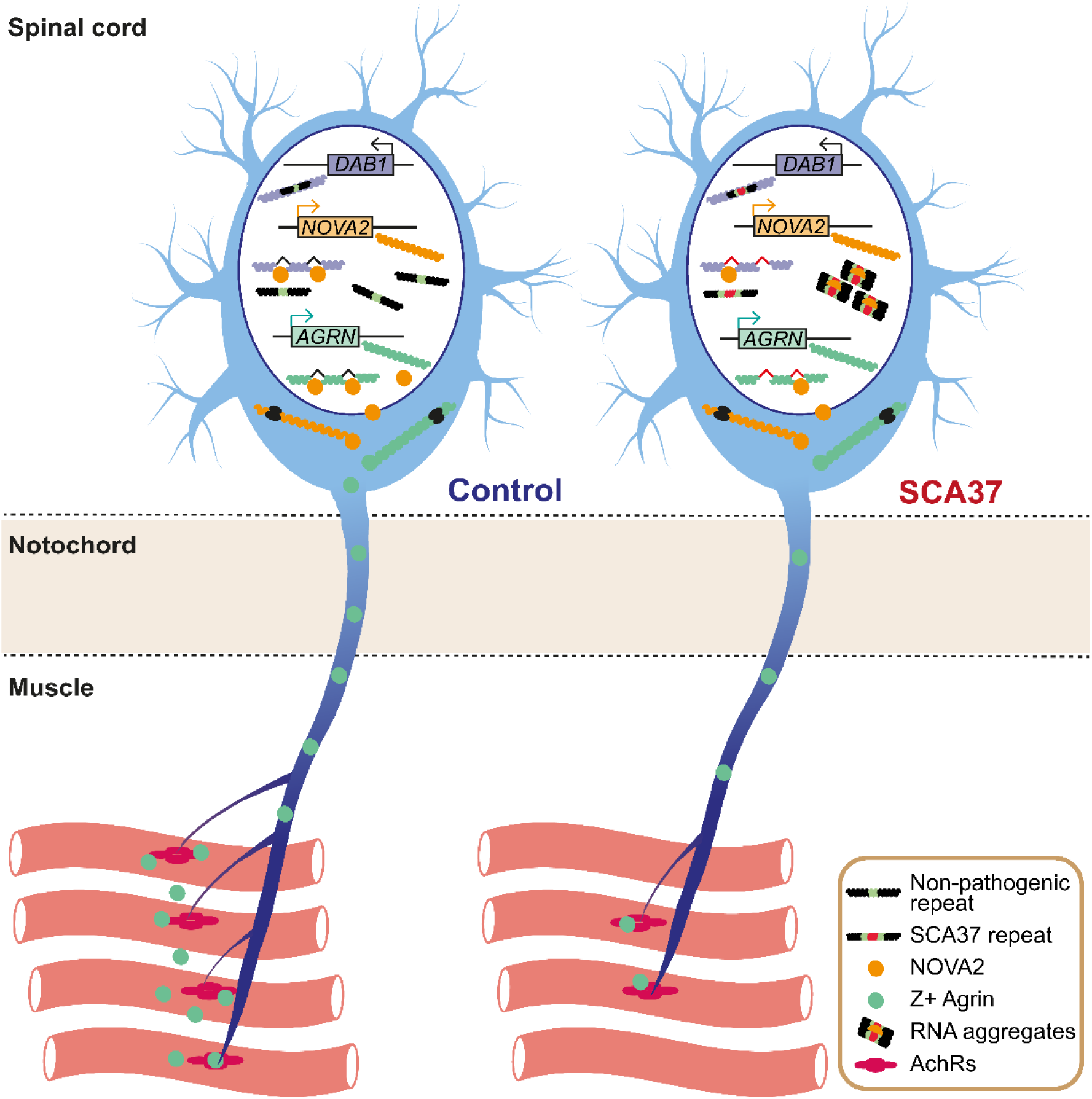
Proposed model describing the neurodevelopmental axonopathy caused by the (AUUUC)_58_ RNA in SCA37. This model illustrates the impact of the (AUUUC)_58_ RNA on spinal cord motor neurons, which disrupts normal axonal outgrowth affecting NMJ formation along muscle cells, as observed in Fig. 3B,C. The stabilization of AchR clusters through altered activity of proteins like NOVA2 and Agrin might also be compromised. Left: During development, *DAB1* with the non-pathogenic repeat is expressed in spinal cord motor neurons, where it plays a vital role in their axonal outgrowth. In these neurons, NMJs are established through the growth of axonal terminals towards prepatterned AchR clusters in muscle tissue. Additionally, NOVA2 is responsible for regulating alternative splicing of neurodevelopmental genes like the neuron-specific Z+ Agrin isoforms, which are transported across neuronal axons and released along muscle tissue, leading to the stabilization of AchRs clusters. Right: In SCA37 motor neurons, the (AUUUC)_58_ RNA, which is transcribed in an intron of *DAB1*, might form aggregates that sequester RBPs like NOVA2, causing its loss-of-function. This NOVA2 dysfunction might lead to abnormal splicing isoforms, leading to the shortening of PMN axons, which was rescued by NOVA2 in Fig. 7C,D. Additionally, the loss-of-function of NOVA2 imprisoned in the (AUUUC)_58_ RNA aggregates results in a decreased Z+ Agrin level, which in turn might prompt the abnormal clustering and mispositioning of AchRs near the horizontal myoseptum seen in Fig. 3B,C.

### The impact of the pathogenic AUUUC repeat RNA beyond SCA37

Pentanucleotide repeat insertions have been described to cause several neurological diseases, as is the case for SCA10, SCA37, and FAMEs (Corbett et al., 2019, Florian et al., 2019, Ishiura et al., 2018, Morato Torres et al., 2022, Seixas et al., 2017, Yeetong et al., 2024, Yeetong et al., 2019). All these diseases are caused by the same or similar repeat motifs and related RNA-mediated mechanisms (Ishiura and Tsuji, 2020, Loureiro et al., 2022, Zhang and Ashizawa, 2022). The seven types of FAME known so far are caused by an ATTTC repeat motif, like SCA37, and are mainly characterized by cortical myoclonic tremors and sporadic tonic-clonic seizures (Corbett et al., 2019, Florian et al., 2019, Ishiura et al., 2018, Yeetong et al., 2024, Yeetong et al., 2019). Notably, there are genes, namely the *CACNA1A*, simultaneously associated with progressive spinocerebellar ataxia and epilepsy (Alonso et al., 2003, Alonso et al., 2004, Stam et al., 2008). The increased number of spontaneous coiling and immobility episodes during development in zebrafish microinjected with the (AUUUC)_58_ RNA might be reminiscent of abnormal excitability. Notably, RNA foci have been detected in cortical and Purkinje cell neurons in FAME1 post-mortem brain tissue (Ishiura et al., 2018), endorsing a common pathogenic mechanism mediated by the (AUUUC)_n_ RNA in FAMEs and SCA37.

### The SCA37 AUUUC repeat RNA causes neurodevelopmental defects

Emerging data, namely from human fetuses, have supported that expression of pathological expanded repeats during embryonic development leads to critical malformations regarding neural cell progenitor differentiation and neuronal connectivity, setting the stage for adult-onset neurodegeneration (Barnat et al., 2020, Hickman et al., 2021, Shabani and Hassan, 2023). Since neurodevelopment is key for the establishment of brain neuronal circuits, and the SCA37 ATTTC repeat is highly expressed as a *DAB1* intronic region in early human embryonic stages (Seixas et al., 2017), the (AUUUC)_58_ RNA has the potential to induce embryonic phenotypes in human fetuses. In this work, the SCA37 (AUUUC)_58_ RNA in zebrafish embryos delayed the hatching process, causing severe morphological abnormalities in embryos. Even though many severely affected infants with other SCAs have been reported (Babovic-Vuksanovic et al., 1998, Miyazaki et al., 1996), for now, they have not been examined in SCA37 families (Corral-Juan et al., 2018, Rosenbohm et al., 2022, Seixas et al., 2017). Consequently, this phenotype is likely an exacerbation of the ubiquitous overexpression of the pathogenic RNA, as also occurs with the expanded myotonic dystrophy type 1 (DM1) CUG repeat in microinjected zebrafish embryos (Todd et al., 2014).

The embryonic expression of the (AUUUC)_58_ RNA during the first hours of development was not sufficient to cause Purkinje cell loss in zebrafish, as described in two SCA37 post-mortem patients (Corral-Juan et al., 2018). The differentiation of Purkinje cells was not significantly affected at 120 hpf, because when Purkinje cell differentiation starts in zebrafish at 72 hpf (Kani et al., 2010), the number of (AUUUC)_58_ RNA copies is likely low due to progressive RNA degradation in the first 24 hpf (Todd et al., 2014). In adult zebrafish, despite the apparently similar Purkinje cell arborization among conditions, the decreased expression of zebrin II-positive Purkinje cells in animals microinjected with both non-pathogenic and pathogenic RNA repeats endorses that the (AUUUC)_58_ RNA is very transient in zebrafish to impact these cells. Notably, zebrin II expression dysregulation has been reported for polyglutamine repeat diseases either in mice or patients with motor dysfunction (Bartelt et al., 2024). Unlike humans, zebrin II-negative Purkinje cells have not been documented in the zebrafish cerebellum so far (Bae et al., 2009).

Axon pathology is a common feature in neurodegenerative diseases, including SCAs, where it precedes cerebellar cell loss (Wilson et al., 2023). Motor neuron involvement with muscle atrophy is present in several SCAs, namely in SCA2, SCA3, and SCA36 (Garcia-Murias et al., 2012, Ghahremani Nezhad et al., 2021, Ikeda et al., 2012, Kanai et al., 2003, Paulson, 2012). The defective axonal outgrowth and muscle cell innervation observed in embryos reflect the disruptive role of the AUUUC repeat RNA in peripheral axons. Similarly, PMN axonal guidance defects in zebrafish embryos modeling SCA13 and C9ORF72 frontotemporal dementia/amyotrophic lateral sclerosis (FTD/ALS) recapitulate the neuronal axonopathy that characterizes ALS (Issa et al., 2012, Swinnen et al., 2018). While developmental synaptic abnormalities in the brain were shown by the defects in climbing fiber innervation of cerebellar Purkinje cells in the SCA1 mouse model expressing an expanded CAG repeat (Ebner et al., 2013); this originates adult-onset motor deficits, which is further supported by the expression of the expanded SCA1 CAG repeat at specific developmental stages (Serra et al., 2006, Zu et al., 2004).

### The embryonic SCA37 (AUUUC)_58_ RNA drives motor dysfunction in adult zebrafish

Aberrant neural circuits have been described for SCAs (Edamakanti et al., 2018, Hsieh et al., 2020, Issa et al., 2012). Our analysis of the NMJ in (AUUUC)_58_ RNA microinjected embryos showed abnormal innervation of muscle cells triggered by abnormal motor neuron outgrowth and, consequently, pathfinding. A similar abnormal innervation pattern is expected in these animals for other neurons in the nervous system. Because precise neuronal circuits need to be established very early in embryonic development, failure to connect properly in these zebrafish animals is expected to cause functional impairment. In fact, adult zebrafish embryonically microinjected with the SCA37 (AUUUC)_58_ RNA presented motor dysfunction, which indicates that embryonic changes in synaptic connectivity can contribute to an adult-onset motor phenotype.

### The SCA37 AUUUC repeat triggers NOVA2 loss-of-function

Transient expression of DM1 and FTD/ALS expanded RNAs in zebrafish has produced developmental phenotypes resulting from RNA-mediated pathogenesis, through RBP sequestration in these late-onset neuromuscular and neurodegenerative disorders (Swinnen et al., 2018, Todd et al., 2014). Since co-expression of human *NOVA2* mRNA with the (AUUUC)_58_ RNA rescued the axonal outgrowth defects in embryos, this indicates a role for NOVA2 loss-of-function in SCA37 pathogenesis. In zebrafish, morpholino-knockdown of *NOVA2* orthologue impaired axon outgrowth, decreasing the number of inter-tecta axonal tracts (Mattioli et al., 2020). In mice, loss-of-function of *Nova2* has also shown defective ventral diaphragm innervation by motor neurons (Saito et al., 2016). Altogether, this evidence strongly suggests that the neurodevelopmental defects are mediated by RBP-impaired function. Indeed, the expression of the SCA37 (ATTTC)_n_ in human cells has shown nuclear RNA aggregation (Seixas et al., 2017). Therefore, we propose that (AUUUC)_58_ RNA aggregates are at the core of the neuronal axon pathology seen in our embryos. In addition, NOVA2 regulates the correct alternative splicing of 540 genes in mice, including *Dab1* (Yano et al., 2010). Moreover, an abnormally spliced *DAB1* isoform has been found in cerebellar tissue from elderly SCA37 patients, not present in non-affected age-matched control individuals, which led the authors to propose embryonic effects of the pathogenic (ATTTC)_n_ insertion as an early trigger of SCA37 pathogenesis (Corral-Juan et al., 2018). NOVA2 regulates the alternative splicing of axon guidance-related genes in mice (Saito et al., 2016), being a key player in synapse formation at the NMJ. It regulates the levels of *Z+ agrin* isoforms released by motor neurons to control the clustering of AchRs in muscle (Ruggiu et al., 2009). This evidence further supports the importance of NOVA2 loss-of-function in the synaptic defects triggered by the SCA37 AUUUC repeat RNA (**Fig.8**).

In conclusion, embryonic SCA37 (AUUUC)_58_ RNA causes presynaptic axonopathy and impaired innervation at the NMJ during development, leading to adult-onset motor dysfunction. This axonopathy is rescued by the NOVA2 RBP, pointing to its role in SCA37 pathogenesis. This knowledge will contribute to delineating approaches for timely therapeutic intervention with significance far beyond SCA37.

## Materials and Methods

### Animal husbandry

Zebrafish (*Danio rerio*) husbandry and procedures were implemented following the National guidelines and legislation for housing and care of laboratory animals (Decreto-Lei n° 113/2013) and the European Union legislation on the protection of animals used for scientific purposes (Directive 2010/63/EU), being authorized by the i3S animal ethics committee (ORBEA - Órgãos Responsáveis pelo Bem-Estar dos Animais, ref. 2018-29) and Direção-Geral de Alimentação e Veterinária (DGAV; Ref. 022872/2020-12-31). Maintenance and handling of zebrafish animals were performed in the i3S zebrafish animal facility, which is licensed by DGAV and part of the AAALAC - International accredited animal care and use program.

### *In vitro* RNA synthesis

The non-pathogenic allele with (ATTTT)_7_, which is one of the most prevalent alleles among the non-affected population, and the pathogenic allele with the (ATTTT)_57_(ATTTC)_58_(ATTTT)_>55_ insertion, hereby designated as (ATTTC)_58_, both alleles with the flanking AluJb element sequences, which contain the right (RM) and the left (LM) monomers, were previously cloned into T7-pCDH-CMV-MCS-EF1-GFP-T2A-Puro vectors and used in our published work (Seixas et al., 2017). To *in vitro* synthesize the (AUUUU)_7_ and (AUUUC)_58_ RNAs, these T7-pCDH-CMV-MCS-EF1-GFP-T2A-Puro DNA vectors were linearized with NotI (Anza) restriction enzyme and purified with DNA Clean & Concentrator kit (ZYMO Research). *In vitro* transcription was performed with T7 RNA polymerase (Thermo Fisher Scientific), and the RNA was then purified with RNA Clean & Concentrator kit (ZYMO Research), aliquoted and stored at -80°C.

### Zebrafish microinjection

To generate embryos for microinjections, adult wild-type strain AB zebrafish animals were crossed, and the collected embryos were microinjected at the 1 to 2-cell stage with approximately 5 nl of (AUUUU)_7_ or (AUUUC)_58_ RNAs at 100 ng/µl, as described in Seixas et al. (Seixas et al., 2017), or purified H_2_O, using the Narishige IM-300 microinjector. As RNAs were diluted in H_2_O, embryos microinjected with this vehicle were used as a control for microinjection toxicity (vehicle condition); as human repetitive RNA can be toxic for zebrafish, animals microinjected with the (AUUUU)_7_ were used as an RNA control from the *DAB1* repeat region in the reference human genome. To control RNA integrity, we monitored the embryo lethality rate and morphological malformations in the (AUUUC)_58_ RNA condition at 100 ng/µl in each microinjection session, according to Seixas et al. (Seixas et al., 2017). A positive toxicity control with the (AUUUC)_58_ RNA at 150 ng/µl was also used in all microinjection sessions, and the replicate experiment was only accepted when this positive control was lethal/toxic for more than 60% of the microinjected embryos. Each independent microinjection session was considered as an experimental replicate. Embryos were maintained at 28.5°C in Petri dishes containing embryo medium (E3) (5 mM NaCl, 0.17 mM KCl, 0.33 mM CaCl2, 0.33 mM MgSO4, 0.00015% methylene blue), supplemented with 0.003% 1-phenyl-2-thiourea (PTU) for imaging purposes, until 144 hours postfertilization (hpf). Embryos developmental stage was assessed by the number of somites, according to Kimmel et al. (Kimmel et al., 1995). After 144 hpf, the microinjected animals were raised under a 14:10h light:dark cycle in the first month in 1 L containers and, then in 3.5 L tanks placed in a water recirculating system with controlled temperature (∼27°C), pH (∼7) and conductivity (∼900 µS). Adult animals were maintained at a maximum density of 8 fish/L in each tank.

### Zebrafish developmental arrest and malformations

For assessing zebrafish developmental arrest and malformations, at least 100 embryos per experimental condition and replicate were microinjected, and blind analysis of embryo morphology was carried out at 24-26 hpf, using a Leica KL 300 LED stereomicroscope (Leica Microsystems). Dead embryos were identified by lack of heartbeat and blood flow, counted, and removed. Embryos were scored according to the severity of abnormalities in head and tail; score 0 for no morphological defects, score 1 for tail defects, score 2 for head and tail defects and score 3 for developmental arrest. To acquire representative images for morphological score analysis, embryos were anesthetized with 0.168 mg/ml tricaine (ethyl 3-aminobenzoate methanesulfonate) and imaged with a 2x, 5x or 8x zoom using a stereomicroscope Leica M205 FA (Leica Microsystems) equipped with an Orca Flash 4.0LT imaging system (C11440-42U30, Hamamatsu Photonics). Number of embryos that survived and hatched was assessed daily from 0 to 96 hpf. At 24 hpf, the scored 0 embryos with no morphological phenotype (undistinguished from the wild-type) were selected for further analyses, hereafter.

### Whole-mounted immunofluorescence

Whole-mounted immunofluorescence was performed at 24 hpf and 120 hpf zebrafish animals. For axon analysis, microinjected embryos at 24 hpf were fixed for 3 h at 4°C in 4% PFA in PBS-T (0.1% Triton X-100 in PBS) and washed three times with PBS-T. For permeabilization and blocking, embryos were incubated in 1% Triton X-100 in PBS for 2 h at RT and then in 1% BSA 1% DMSO 1% NGS 0.7% Triton X-100 in PBS for 1 h at RT. Then, samples were incubated with anti-synaptic vesicle 2 (SV2) antibody (1:200, mouse, AB2315387, Developmental Studies Hybridoma Bank (DSHB)), diluted in blocking solution ON at 4°C. Then, embryos were washed three times with PBS-T and incubated with the secondary antibody Alexa Fluor-647 goat anti-mouse IgG (1:750, A-21235, Thermo Fischer Scientific) and DAPI (1:1000, Sigma-Aldrich, Merck), diluted in blocking solution for 4 h at RT. For the neuromuscular junction (NMJ) analysis, embryos were also incubated with the postsynaptic acetylcholine receptor (AchR) marker, tetramethylrhodamine conjugated α-Bungarotoxin (α-BTX, 1:1000, T1175, Thermo Fischer Scientific), diluted in blocking solution for 4 h at RT. Embryos were washed three times with PBS-T and mounted on microscope slides using glycerol-based ibidi mounting media (50011, ibidi). For zebrin II-positive Purkinje cells analysis, larvae were fixed at 120 hpf and incubated with an anti-zebrin II antibody (1:50, kindly gifted by Dr Richard B. Hawkes) (Lannoo et al., 1991a, Lannoo et al., 1991b) and the secondary antibody Alexa Fluor-647 goat anti-mouse IgG (1:750, A-21235, Thermo Fischer Scientific), in the same conditions described above. Images were acquired on an inverted Leica SP8 single point scanning confocal microscope equipped with a fully motorized DMi8 microscope (Leica Microsystems) a HC PL APO 40x /1.10 Water CS2 (with motorized correction ring) objective lens (see **Supplementary Materials and Methods**). For zebrin II-positive Purkinje cell analysis, the same system was used with a 25x objective (NA 0.95; see **Supplementary Materials and Methods**). To analyze axonal length of primary motor neurons (PMNs), the 6 axons in the 6-somites spanning region immediately anterior to the cloaca were measured to guarantee the analysis of the same axons in all conditions; an average intensity z-projection of a minimum of five zebrafish microinjected embryos for each experimental condition and replicate was used to perform the axon tracing using the Fiji plugin NeuronJ (Meijering et al., 2004, Schindelin et al., 2012). Briefly, the axon tracing was performed manually from the closest point to the neuron soma to the end of the axonal terminal, and the average axonal length was calculated for each embryo. For co-localization analysis of presynaptic SV2 and postsynaptic α-BTX markers, images of at least six previously microinjected embryos from each condition and replicate were deconvolved using the Deconvolution Express module and the Standard algorithm from Huygens Professional, followed by the calculation of Pearson’s co-localization coefficient (version 20.04, SVI). The AchRs distribution in the dorsal-ventral axis of muscle in zebrafish embryos was assessed in z-projections (maximum intensity) using the Fiji plugin NeuronJ (Meijering et al., 2004, Schindelin et al., 2012), starting the measurement in the horizontal myoseptum for both dorsal and ventral orientations. The number of AchRs by means of α-BTX staining was determined by using the Object Analyzer tool from Huygens Professional and the SV2 staining as a reference for segmentation (version 20.04, SVI). To determine the area of AchRs, a z-maximum intensity projection was made and used to manually define the regions of interest based on the α-BTX staining. For the analysis of Purkinje cells, IMARIS software (version 10.2.0, Andor Oxford) was used to determine the volume and the mean intensity of zebrin II-positive cells in at least nine larvae from each condition per replicate.

### Spontaneous tail coiling assay

For zebrafish spontaneous tail coiling assessment at 24 hpf, the microinjected embryos were placed in a Petri dish and acclimatized to the room light for 30 minutes. The number of tail movements was counted per embryo for 60 seconds, blindly to the embryo conditions, under a Leica KL 300 LED stereomicroscope. Representative videos of spontaneous coiling were acquired using a stereomicroscope Leica M205 FA (Leica Microsystems) equipped with an Orca Flash 4.0 LT imaging system (model C11440-42U30, Hamamatsu Photonics) controlled using the Micro-Manager FIJI plugin (Edelstein et al., 2010).

### Touch-evoked escape swimming response test

To test the escape response to a tactile stimulus at 72 hpf, at least 20 larvae were blindly tested per condition and in replicates (a total of 6 replicates). For acclimatization, at 24 hpf, the microinjected embryos were placed in a room with controlled temperature (∼27°C) under a 14:10h light:dark cycle. The animals were acclimatized to the intensity of the light box for at least 30 minutes before the test started at 72 hpf. Then, each larva was placed in a Petri dish on top of a light box with a white acrylic panel to reduce light intensity. Once the free-swimming movement stopped, each larva was gently touched on the tail with a black filament. Swimming responses were recorded using a GoPro Hero8 Black (C3331350868456) at 120 frames per second until the larva stopped swimming. Touch-evoked escape response was assessed as no response or presence of response (any movement or swimming response). Also, the time of reaction between the touch and the start of the response, and the swimming duration (time that larvae swim after touch) were analyzed. The swimming response is characterized by a movement away from the touch-escape response. The time of reaction and swimming duration were analyzed, frame by frame, using Fiji software (Schindelin et al., 2012). Larvae that did not show any response were excluded from the analyses of the time of reaction and the swimming duration; larvae that swam to the border of the Petri dish were excluded from the swimming duration analysis.

### Spontaneous locomotor behavior test

To assess spontaneous motor activity, the spontaneous locomotor activity test was performed in 20 larvae per condition and replicate at 144 hpf, in 6-well plates (3 replicates). For each well, 5 ml of 0.5% agarose diluted in the fish-recirculating water system was polymerized, and a circular hole (radius ∼1.9 mm) was made in the center of each agarose well. The wells were filled with 2 ml of water from the fish recirculating system. Each larva from the conditions studied was blinded anonymized and randomized per well. Larvae locomotor activity was recorded with a GoPro Hero8 Black (C3331350868456) at 60 frames per second for 10 minutes, using a zoom of 2. Larvae tracking was performed using Any-Maze software (Stöelting, Dublin, Ireland). The locomotor activity was assessed by percentage of immobility in the 10 min, swimming distance (cm), average speed (cm/s), time immobile (s) and number of immobile episodes. Immobility was considered when no movement was detected for more than 2 seconds. If correct tracking was not possible, those movies were discarded.

### Novel diving tank test

The novel diving tank test was performed in 3- and 17-month-old animals to assess locomotor activity in adult fish in a novel environment. Microinjected embryos without morphologic malformations from five independent microinjection sessions were raised. Approximately 25 adult fish per condition were placed in 3.5 L tanks. For the novel diving tank test, each fish was placed in a 3.5 L tank filled with 2.5 L of fresh water (12 cm of water column) from the fish recirculating system, and its locomotor activity was recorded at 25 frames per second for 6 minutes by a side-view camera (Canon Legria HF R606). The videos were converted from MP4 to AVI format using Any Video Converter software and fish tracking was performed using the open-source software TheRealFishTracker (Buske and Gerlai, 2014). The percentage of immobility, swimming distance (m) and average speeds (cm/s) were calculated for each fish using the coordinates given by the software. After the test, animals were euthanized with 0.300 g/L of tricaine, and standard length and weight were measured in each fish. Brains were collected for further histological analysis.

### Zebrafish brain histology and immunofluorescence

Zebrafish brains were dissected and fixed in 4% PFA in PBS ON at 4°C, and then, they were washed 3 times with PBT (PBS with 0.1% Tween) and stored in methanol at -20°C. Tissues were preprocessed by dehydration through an increasing series of ethanol concentrations, clearance with xylene and a final step with paraffin, for 10 minutes in each reagent. Brains were embedded in paraffin, and 5 µm slices were longitudinally sectioned using a Leica RM2255 microtome (Leica Biosystems). Before staining and immunofluorescence, deparaffinization and rehydration of slices were performed in the reverse order compared to the previous protocol. Then, slices were stained with hematoxylin and eosin or underwent an anti-zebrin II antibody immunofluorescence protocol. For immunofluorescence, slices were submitted to antigen retrieval with 10 mM EDTA pH 8.0 in a steamer for 40 min, followed by 20 min at RT. Immunofluorescence protocol with zebrin II antibody (1:50, kindly gifted by Dr Richard B. Hawkes) (Lannoo et al., 1991a, Lannoo et al., 1991b) was performed as described above. Imaging was performed using the laser spinning disk confocal microscope Andor BC43 (Andor Oxford Instruments), equipped with a CFI Plan Apochromat Lambda D 20X/0.8 DIC (see **Supplementary Materials and Methods**); a minimum of three animals were analyzed per condition. Hematoxylin and eosin-stained slices were digitized using the Phenoimager^TM^ HT (Akoya Biosciences), and the area of the cerebellar molecular cell layer was measured in ten consecutive slices for each brain using the QuPath-0.5.1 (Bankhead et al., 2017). The average number of zebrin II-positive Purkinje cells was quantified in five consecutive slices of each brain using anti-zebrin II fluorescence in IMARIS software (version 10.2.0, Andor Oxford Instruments).

### RBP *in silico* analysis

To identify potential RBPs that bind to the non-pathogenic and SCA37 (AUUUC)_58_ RNAs, the RBPmap webserver was used (Paz et al., 2014). Given the complete length of the pathogenic repeat allele of approximately 170 repeat units (Seixas et al., 2017), the non-pathogenic (AUUUU)_170_ was chosen to be introduced in this website, as well as the complete repeat sequence of the pathogenic (AUUUU)_57_(AUUUC)_58_(AUUUU)_>55_, which corresponds to its abbreviated name (AUUUC)_58_ RNA; all human/mouse RBP motifs from the database were selected, and default stringency and conservation filter were applied. From the z-scores of each predicted binding site (p<0.05), the maximum z-score was determined for each RBP hit. The number of putative binding sites of each RBP was calculated for non-pathogenic and SCA37 (AUUUC)_58_ RNAs. This analysis was performed to identify differences in the binding affinity of each RBP between non-pathogenic and SCA37 (AUUUC)_58_ RNAs.

### NOVA2 experiments in zebrafish

To synthesize human *NOVA2* mRNA for microinjection in zebrafish, a sequence encoding fourteen histidine residues was cloned in pCMV6-XL6-NOVA2 (SC303210, Origene) at the C-terminal of NOVA2 cDNA using the IVA cloning approach (Garcia-Nafria et al., 2016), for NOVA2 immunofluorescence staining with anti-histidine antibody. Amplification of *NOVA2* was performed using Phusion High Fidelity DNA polymerase (NEB) with the following primers: 5’-CATCATCATCATCATCATTGAGGCCTGTGGTGTGTGCTC-3’ and 5’-ATGATGATGATGATGATGTCCCACTTTCTGGGGGTTTGAGG-3’. Human *NOVA2* mRNA was *in vitro* synthesized from the pCMV6-XL6-NOVA2-Histag DNA vector. After DNA plasmid linearization with XbaI (Anza) restriction enzyme and purification with DNA Clean & Concentrator kit (ZYMO Research), *NOVA2* cDNA was *in vitro* transcribed with SP6 RNA polymerase (Thermo Fisher Scientific), and the mRNA was purified with RNA Clean & Concentrator kit (ZYMO Research), aliquoted and stored at -80°C. Zebrafish embryos were microinjected and maintained as described above. Translation of the human NOVA2 protein in zebrafish was confirmed by whole-mount immunofluorescence with anti-histidine-tagged antibody (1:500, 05-949, Milipore) in 24 hpf zebrafish embryos. Briefly, embryos were fixed for 3 h at 4°C in 4% PFA in PBS-T, washed three times with PBS-T, and permeabilized with 1% Triton X-100 in PBS for 2 h at RT. Samples were blocked with 5% BSA in PBS-T for 1 h at RT and incubated with anti-histidine-tagged antibody (1:500, 05-949, Milipore) diluted in blocking solution ON at 4°C. Then, embryos were washed three times with PBS-T and incubated with the secondary antibody Alexa Fluor-647 goat anti-mouse IgG (1:750, A-21235, Thermo Fischer Scientific) and DAPI (1:1000, Sigma-Aldrich, Merck) diluted in blocking solution for 4 h at RT. Whole-mount immunofluorescence with SV2 antibody (1:200, mouse, AB2315387, Developmental Studies Hybridoma Bank (DSHB)) and analysis were also performed in 24 hpf embryos, as described above. Images were acquired on an inverted Leica SP8 single point scanning confocal microscope equipped with a fully motorized DMi8 microscope (Leica Microsystems) a HC PL APO 40x /1.10 Water CS2 (with motorized correction ring) objective lens (see **Supplementary Materials and Methods**).

### Statistical analysis

GraphPad Prism was used for the graphical representation of the data, and statistical analysis was performed using IBM SPSS. Shapiro-Wilk and Levene’s tests were used to assess the normality of the data and homogeneity of variances, respectively. Data were transformed (logarithmic or cube root transformation) when not normally distributed. If transformed data are not normal, non-parametric tests were used. The ꭕ^2^ test was used for n≥50 to compare categorical variables. To evaluate differences among experimental groups, the one-way analysis of variance (ANOVA) following the Bonferroni post-hoc test was used for normally distributed data, and the Kruskal-Wallis test following Dunn’s test as a post-hoc test was used for non-normal data. When only the homogeneity of variances of the data is not verified, Welch’s correction was applied to the one-way ANOVA, and Dunnett’s T3 post-hoc test was used. Two-way ANOVA was used to study how two independent variables (e.g. condition and age or condition and replicates) affect a dependent variable. For comparisons between two independent samples, the independent Student’s t-test or the Mann-Whitney U Test was used for normal or non-normally distributed data, respectively. The Log rank test was used for zebrafish survival and hatching rates. Percentage or mean values and standard deviation details are described in the results section. The sample size and replicates relative to each analysis are indicated in the legend of each figure. Significant differences were considered when p≤0.05 (two-tailed).

## Acknowledgements

We thank Prof. Richard B. Hawkes for the anti-zebrin II antibody; zebrafish i3S animal facility staff (associated with the Vertebrate Development and Regeneration group, José Bessa), including Joana Marques and Isabel Guedes for maintenance of the zebrafish; Dr. Tiago Silva for imaging analyses insights; Dr. Sara Ricardo for her expertise and support with antigen retrieval procedure and, i3S Scientific Platforms Advanced Light Microscopy and Histology and Electron Microscopy, members of the national infrastructure PPBI-Portuguese Platform of BioImaging (supported by POCI-01-0145-FEDER-022122) for the support with microscopy, imaging and histological analyses.

## Funding

This work was funded by National Funds through FCT—Fundação para a Ciência e a Tecnologia, I.P., under the project UIDB/04293/2020 POCI-01-0145-FEDER-029255 (PTDC/MED-GEN/29255/2017); by COMPETE2030-FEDER-00691000, Grant No. 15801, funded by COMPETE2030 and FCT under MPr-2023-12; and FCT PhD scholarships (2020.00528.BD and 2021.05757.BD).

### Data and resource availability

All relevant data and resources can be found within the article and its supplementary information.

### Authors contributions

Conceptualization: A.F.C, A.M.V., J.B., I.S.

Methodology: A.F.C., A.S.F., J.R.L., M.M.A., P.S., A.M.V., J.B., I.S.

Investigation: A.F.C., A.S.F. Formal analysis: A.F.C., A.M.V.

Writing - original draft: A.F.C., J.B., I.S.

Writing – review & editing: A.F.C., A.S.F., J.R.L., M.M.A., P.S., A.M.V., J.B., I.S.

All authors read and approved the final manuscript.

