## Supplemental Information for "Embryonic Spinocerebellar Ataxia Type 37 AUUUC Repeat RNA Causes Neurodevelopmental Defects in Zebrafish"

*Equal contribution

•Corresponding author:

**Isabel Silveira, PhD**

Genetics of Cognitive Dysfunction Laboratory, i3S - Instituto de Investigação e Inovação em Saúde and IBMC - Institute for Molecular and Cell Biology, University of Porto, Portugal, Rua Alfredo Allen, 208, 4200-135 Porto, Portugal

**José Bessa, PhD**

Vertebrate Development and Regeneration Laboratory, i3S - Instituto de Investigação e Inovação em Saúde and IBMC - Institute for Molecular and Cell Biology, University of Porto, Portugal, Rua Alfredo Allen, 208, 4200-135 Porto, Portugal

### Supplementary Materials and Methods

**Embryo/larva imaging on inverted Leica SP8 single point scanning confocal microscope**

Zebrafish PMNs images were acquired with a 405 Diode laser line, a tunable Argon laser AOBS modulated and 633 HeNe laser line, and Leica PMT and HyD detectors. PMNs were imaged sequentially by line scanning unidirectionally at 400 Hz using the galvanometer-based imaging mode and line average of 3, with a pixel size of 0.28 μm and a z step-size of 0.42 μm, for an area size of 290.62 μm x 290.62 μm in Leica LasX software (version 3.5.6.21594) and saved as LIF files. For all images the pinhole size was 77.2 μm, calculated at 1 AU for 580 nm emission. DAPI was excited with 405 nm laser line with laser power of 2.99%, and emission was collected on a Leica PMT2 detector with a collection window of 415 – 478 nm with 601.6 V and an offset of 0. AlexaFluor 647 was excited with 633 nm laser line with laser power of 0.10%, and emission was collected on a Leica HyD4 (standard mode) detector with a collection window of 643 – 743 nm and smart gain of 19.4%. Zebrafish NMJs were imaged sequentially by line scanning bidirectionally at 400 Hz using the galvanometer-based imaging mode and line average of 5, with a pixel size of 0.19 μm and a z step-size of 0.42 μm, for an area size of 193.75 μm x 193.75 μm in Leica LasX software (version 3.5.6.21594) and saved as LIF files. DAPI and AlexaFluor 647 were excited as described for PMNs. Tetramethylrhodamine was excited with 514 nm laser line with laser power of 0.50% (Argon at 29.31%), and emission was collected on a Leica HyD3 detector with a collection window of 524 – 625 nm with a smart gain of 51.6%. Purkinje cell images of 120 hpf larvae were acquired sequentially by line scanning unidirectionally at 400 Hz using the galvanometer-based imaging mode and line average of 3, with a pixel size of 0.38 μm and a z step-size of 0.57 μm, for an area size of 387.5 μm x 387.5 μm in Leica LasX software (version 3.5.6.21594) and saved as LIF files. For all images the pinhole size was 55.7 μm, calculated at 1 AU for 580 nm emission. DAPI was excited with 405 nm laser line with laser power of 1.50%, and emission was collected on a Leica PMT2 detector with a collection window of 415– 478 nm with 600 V and an offset of 0. AlexaFluor 647 was excited with 633 nm laser line with laser power of 0.10%, and emission was collected on a Leica HyD3 detector with a collection window of 643 – 696 nm and smart gain of 50%. To assess human NOVA2 protein translation in zebrafish, images were acquired sequentially by line scanning bidirectionally at 400 Hz using the galvanometer-based imaging mode and line average of 3, with a pixel size of 0.03 μm and a z step-size of 0.42 μm, for an area size of 29.06 μm x 29.06 μm in Leica LasX software (version 3.5.6.21594) and saved as LIF files. For all images the pinhole size was 77.2 µm, calculated at 1 AU for 580 nm emission. DAPI was excited with 405 nm laser line with laser power of 2.99%, and emission was collected on a Leica PMT2 detector with a collection window of 415 – 478 nm with 601.6 V and an offset of 0. AlexaFluor 647 was excited with 633 nm laser line with laser power of 0.10%, and emission was collected on a Leica HyD4 (standard mode) detector with a collection window of 643 – 743 nm and smart gain of 19.4%. In NOVA2 rescue experiments, images were acquired with a 405 Diode laser line and 633 HeNe laser line, and Leica PMT and HyD detectors. PMNs were imaged sequentially by line scanning bidirectionally at 400 Hz using the galvanometer-based imaging mode and line average of 3, with a pixel size of 0.28 µm and a z step-size of 0.42 µm, for an area size of 290.62 µm x 290.62 µm in Leica LasX software (version 3.5.6.21594) and saved as LIF files. For all images the pinhole size was 77.2 µm, calculated at 1 AU for 580 nm emission. DAPI was excited with 405 nm laser line with laser power of 4.09%, and emission was collected on a Leica PMT2 detector with a collection window of 415 – 478 nm with 601.6 V and an offset of 0. AlexaFluor 647 was excited with 633 nm laser line with laser power of 0.30%, and emission was collected on a Leica HyD4 (standard mode) detector with a collection window of 643 – 743 nm and smart gain of 19.4%.

**Adult brain slices imaging on laser spinning disk confocal microscope Andor BC43**

DAPI was excited with 405 nm laser line with laser power of 100%, with an exposure time of 50 ms, and emission was collected with a 445/20 filter. AlexaFluor 647 was excited with 638 nm laser line with laser power of 90%, with an exposure time of 50 ms, and emission was collected by a 708/75 filter.

### Supplementary Figures and Table

A

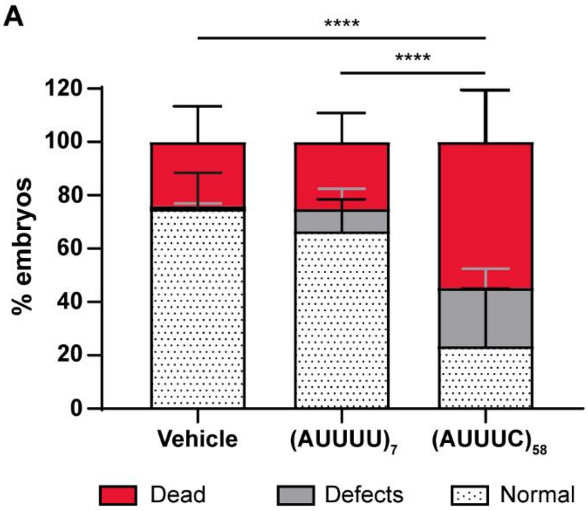


C

| Condition \ Score | 0 | 1 | 2 | 3 |
|---|---|---|---|---|
| Vehicle | 98.81 | 0.38 | 0.67 | 0.14 |
| (AUUUU)$_7$ | 89.53 | 7.90 | 2.39 | 0.18 |
| (AUUUC)$_{58}$ | 47.22 | 24.63 | 26.99 | 1.16 |

B

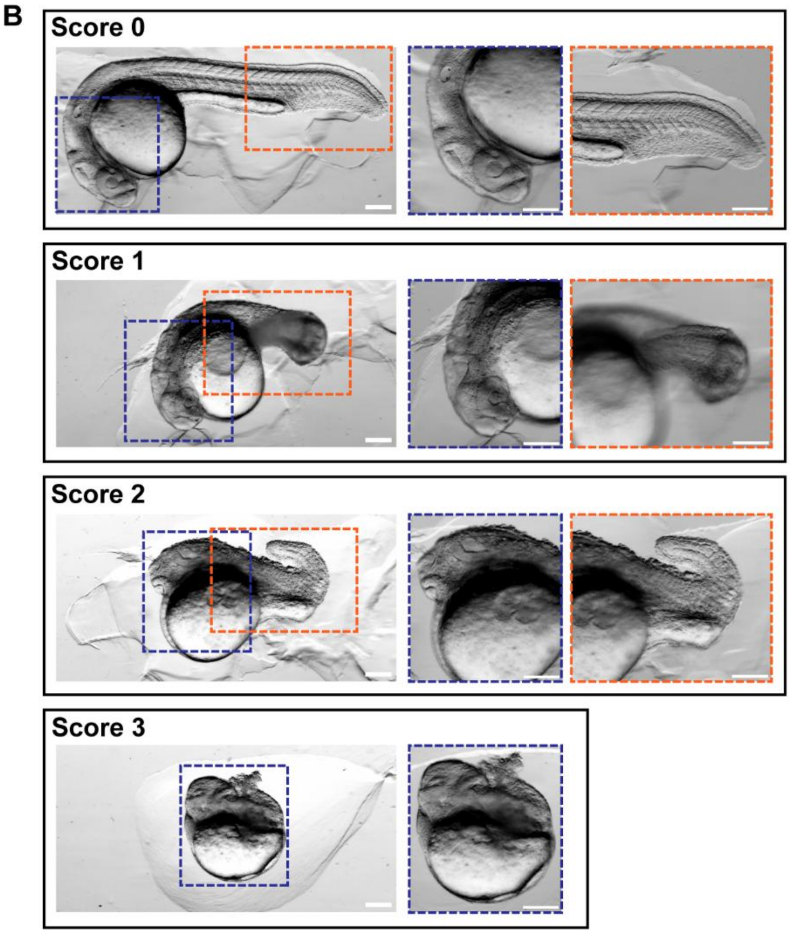


**Fig. S1 The (AUUUC)$_n$ insertion RNA causes developmental toxicity, resulting in arrested development and lethality.** (A) Percentage of dead (red), defective (gray) and normal (white with black dots) embryos observed at 24 hpf following the microinjection of vehicle, (AUUUU)$_7$ or (AUUUC)$_{58}$ RNAs (average of eight independent experiments with at least 100 embryos per experiment; ****$p<0.0001$, $\chi^2$ test for lethality and morphological defects). Data are shown as mean ± standard deviation. (B) Representative images correspond to the phenotypical scores; 0 - embryos without morphological defects, 1 - embryos with defects in tail, 2 - embryos with defects in tail and head, and 3 - arrested embryonic development; respective zoom-in images are presented for blue and orange delimitations; scale bar = 50µm. (C) Percentage of embryos exhibiting different phenotypical scores.

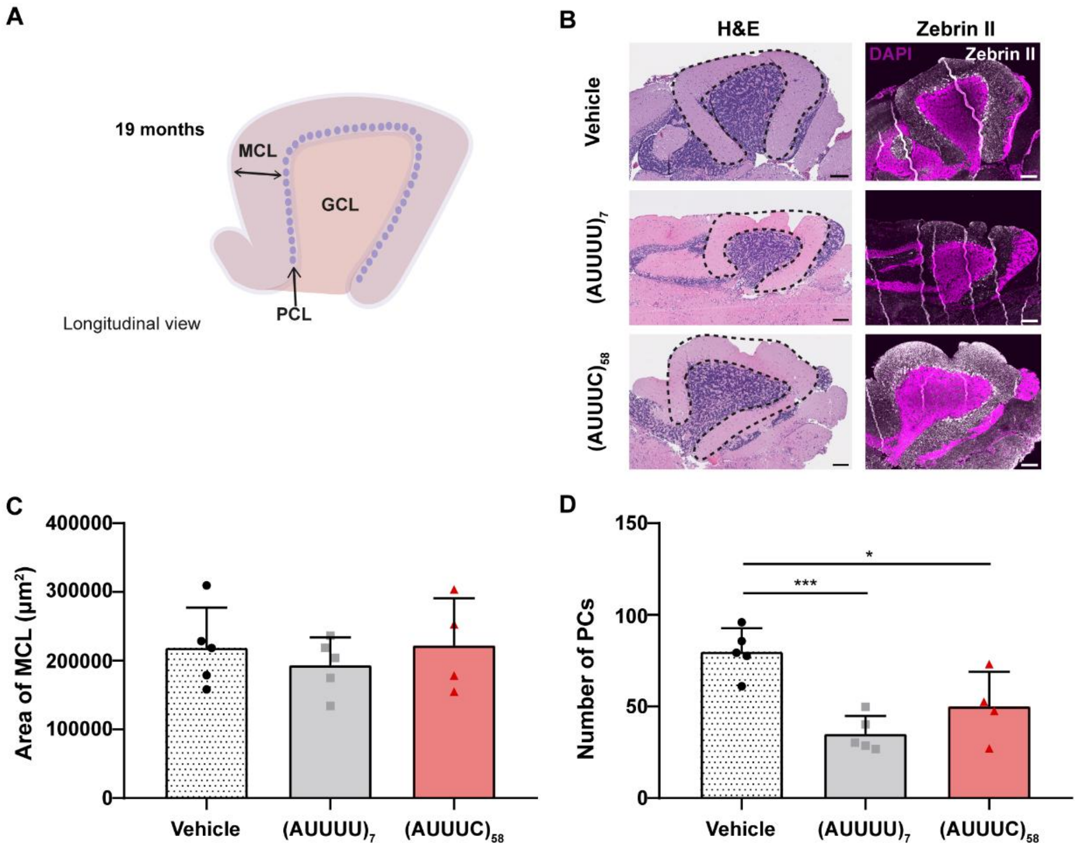


**Fig. S2 No evidence of zebrin II-positive Purkinje cell loss in adult animals previously microinjected with the $(AUUUC)_{58}$ RNA.** (A) The schematic representation illustrates the adult zebrafish cerebellum from a longitudinal view, highlighting the anatomical layout and the location of Purkinje cells within the cerebellar structure. This representation serves as a foundational overview for comparing Purkinje cell presence and distribution in RNA microinjected versus control animals. (B) Representative images depict paraffin brain sections of 19-month-old zebrafish. The left panel shows hematoxylin and eosin staining, providing a general view of the cellular architecture within the cerebellum; all the cerebellar molecular cell layer delimited in black dashed line was measured, independently of brain size. The right panel displays immunofluorescence results using an anti-zebrin II antibody, counterstained with DAPI to highlight cell nuclei. Scale bar = 100µm. (C) The area of the cerebellar molecular cell layer was assessed with sample sizes consisting of n=5 animals in the vehicle group, n=5 animals in the $(AUUUU)_7$ group, and n=4 animals in the $(AUUUC)_{58}$ group. Statistical analysis via One-way ANOVA indicated no significant differences in the area among the conditions. (D) The number of Purkinje cells was quantified across the different animal groups, with n=5 animals in the vehicle group, n=5 animals in the $(AUUUU)_7$ group, and n=4 animals in the $(AUUUC)_{58}$ group. The analysis revealed a significant difference in Purkinje cell number, indicated by a p-value of *p<0.05 and ***p<0.001, as determined by One-way ANOVA followed by Bonferroni correction. Data are shown as mean ± standard deviation. Abbreviations: PC – Purkinje cells; GCL – granular cell layer; PCL – Purkinje cell layer; MCL – molecular cell layer.

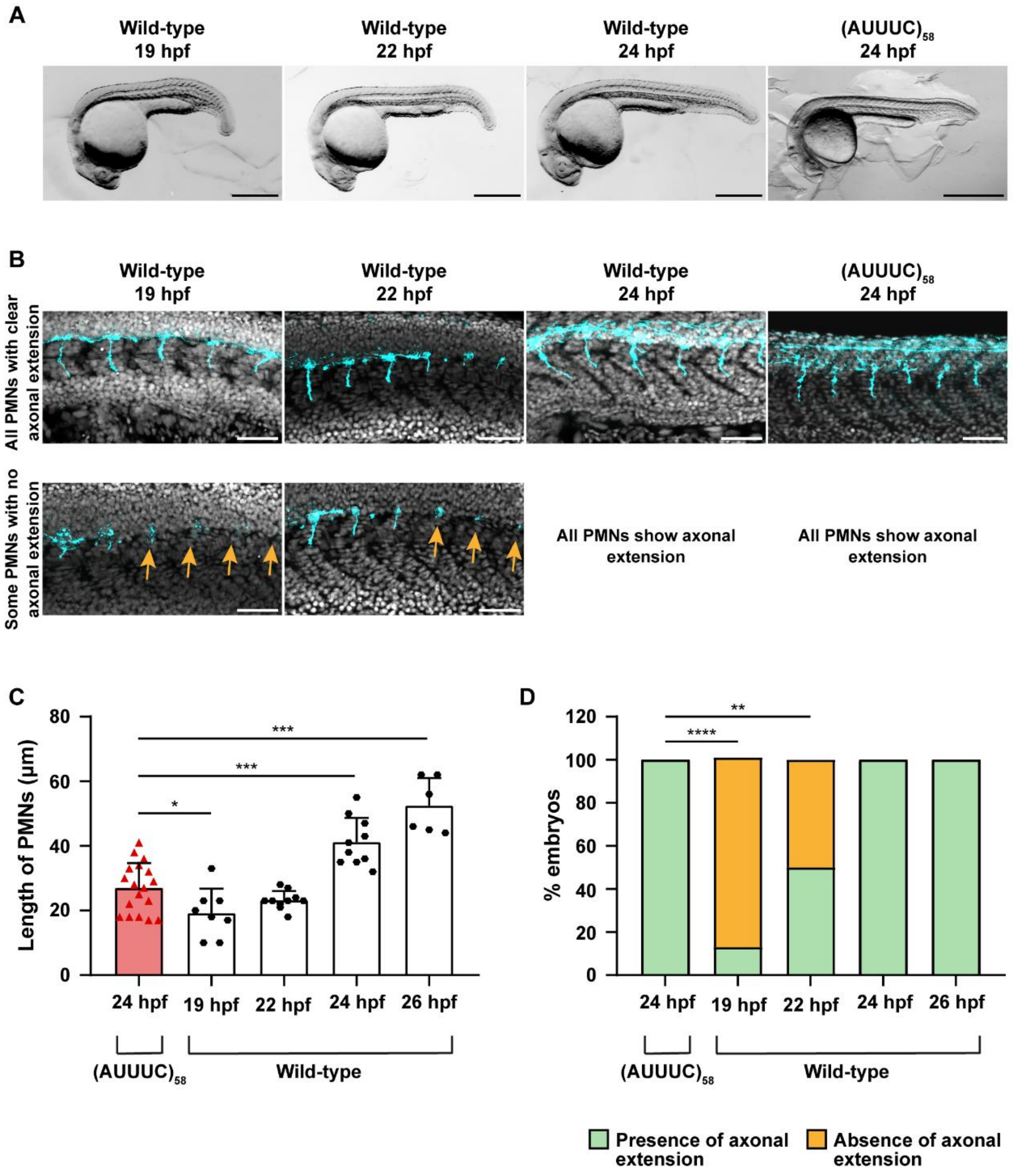


**Fig. S3 The (AUUUC)$_{58}$-injected embryos show axonal extension of PMNs at 24 hpf.** Related to **Fig. 3B.** (A) Representative images showing the morphology of non-injected wild-type embryos at 19, 22 and 24 hpf and (AUUUC)$_{58}$-injected embryos at 24 hpf; microinjected embryos at 24 hpf used for PMN axonal length quantification; scale bar = 200 µm. (B) Representative z-average projection images of PMNs axons from non-injected wild-type embryos at 19, 22 and 24 hpf and from (AUUUC)$_{58}$-injected embryos at 24 hpf, in the 6-somites region anterior to the cloaca (except for 19 hpf, as only one out of eight embryos showed extension of PMNs; scale bar = 50 µm), demonstrating that some PMNs do not present axonal outgrowth at 19 and 22 hpf (orange arrows), contrarily to 24 hpf. (C) Quantification of PMN axonal length in both $(AUUUC)_{58}$-injected and non-injected embryos at different timepoints. Statistical analyses showed significant differences between axonal length of PMNs in $(AUUUC)_{58}$ RNA microinjected embryos at 24 hpf and non-injected embryos at 19, 24 or 26 hpf (n=18 embryos microinjected with the $(AUUUC)_{58}$ RNA, n=8 wild-type embryos at 19 hpf, n=10 wild-type embryos at 22 hpf, n=10 wild-type embryos at 24 hpf, n=6 wild-type embryos at 26 hpf; *$p<0.05$ and ***$p<0.001$, independent Student's t-test). Data are represented as mean ± standard deviation. (D) Percentage of embryos presenting axonal extension of PMNs in the 6-somites region anterior to the cloaca at different timepoints. All the embryos microinjected with the $(AUUUC)_{58}$ RNA at 24 hpf show axonal extension, when compared with the non-injected wild-type embryos at 19 and 22 hpf. Statistical analysis demonstrated significant differences in terms of presence or absence of axonal extension between $(AUUUC)_{58}$-injected embryos and wild-type embryos at 19 or 22 hpf (n=18 embryos microinjected with the $(AUUUC)_{58}$ RNA, n=8 wild-type embryos at 19 hpf, n=10 wild-type embryos at 22 hpf, n=10 wild-type embryos at 24 hpf, n=6 wild-type embryos at 26 hpf; **$p<0.01$ and ****$p<0.0001$, Fisher exact test for number of embryos with or without PMN axonal extension. Data represented as mean.

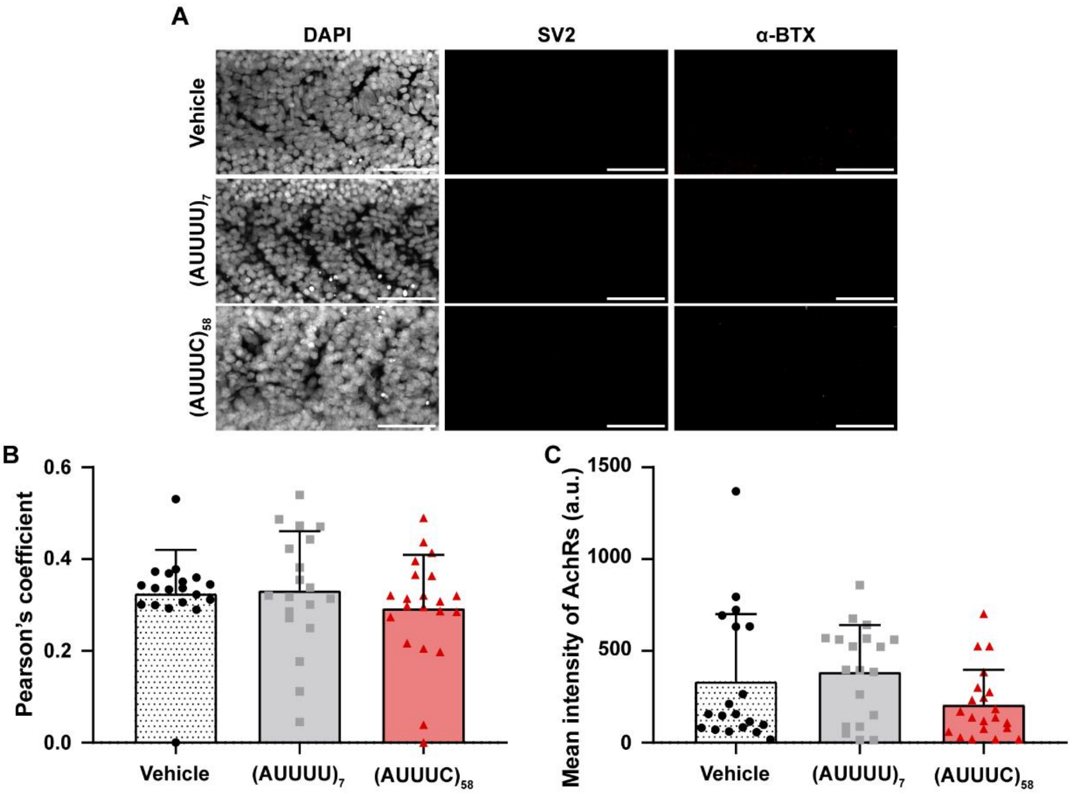


**Fig. S4 NMJs in (AUUUC)$_{58}$-injected zebrafish embryos.** Related to **Fig. 3C.** (A) Representative z-maximum intensity projections showing immunofluorescence without primary antibodies in 24 hpf embryos microinjected with the vehicle, (AUUUU)$_7$ or (AUUUC)$_{58}$ RNAs; scale bar = 50 µm. (B) Co-localization of the presynaptic SV2 and postsynaptic α-BTX markers measured by Pearson's co-localization coefficient, with values ranging from 0 to 1. Data are presented for n=19 embryos in the vehicle and (AUUUU)$_7$ groups, and n=21 embryos in the (AUUUC)$_{58}$ group, sourced from 3 experimental replicates. Statistical analysis was conducted using the Kruskal-Wallis test for the co-localization coefficient, and the results are indicated as not significant. (C) α-BTX mean intensity calculation for NMJs with data collected from n=19 embryos in the vehicle group, n=19 embryos in the (AUUUU)$_7$ group, and n=21 embryos in the (AUUUC)$_{58}$ group, across 3 experimental replicates. Statistical analysis using the Kruskal-Wallis test to assess differences in mean intensity, with results indicated as not significant. Data are represented as mean ± standard deviation.

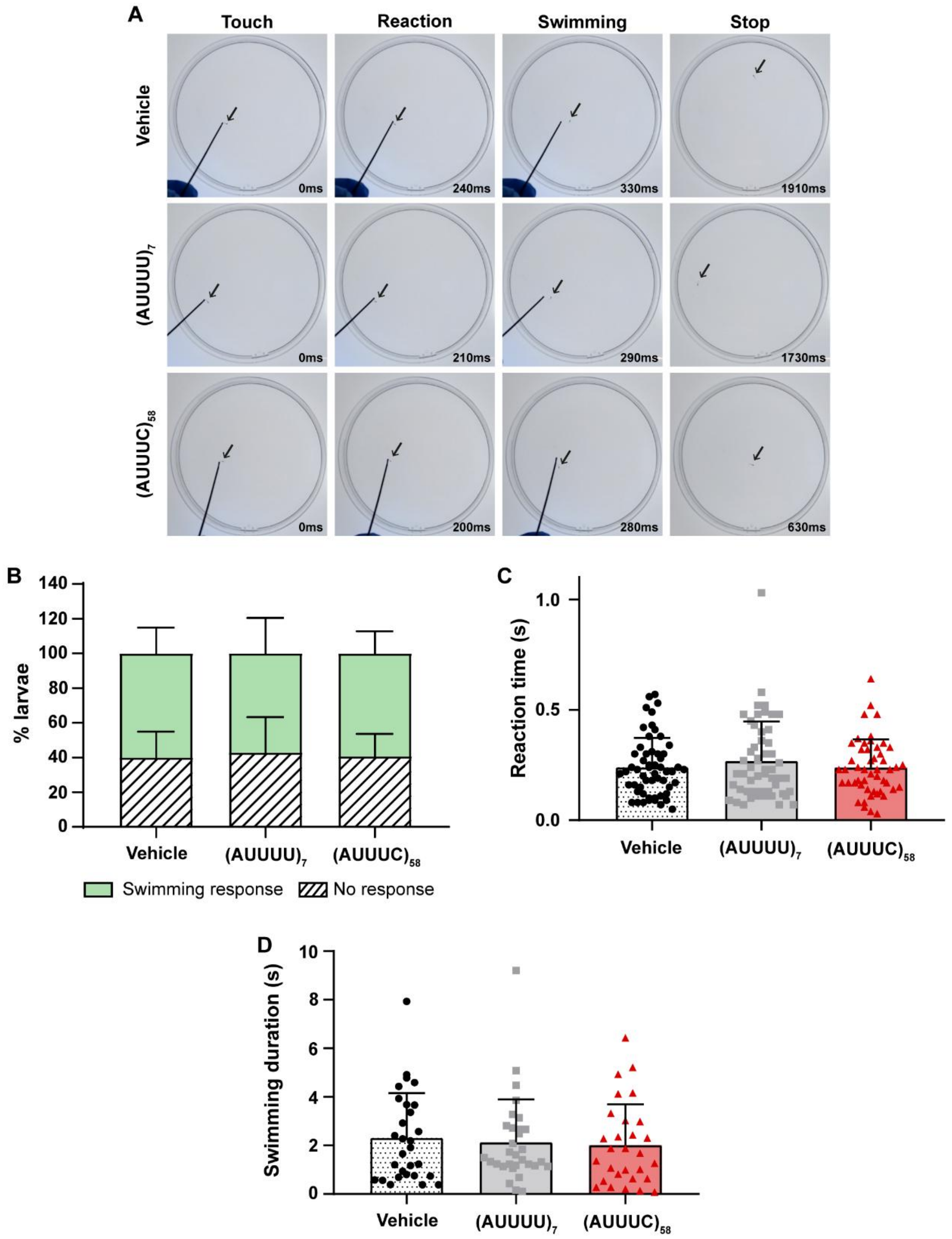


**Fig. S5 (AUUUC)58 larvae exhibited no motor or sensorial defects.** (A) Representative frames showcasing the touch stimuli, the larvae reaction, the swimming away from the filament (10 frames after their reaction) and the cessation of swimming observed during the touch-evoked escape swimming test. This test was performed on larvae microinjected with either the vehicle, (AUUUU)$_7$, or (AUUUC)$_{58}$ at 72 hpf. (B) The data indicates the percentage of larvae that responded or did not respond to the touch stimulus across three different groups: the vehicle group (n=91 larvae), the $(AUUUU)_7$ group (n=88 larvae), and the $(AUUUC)_{58}$ group (n=84 larvae), with a total of six experimental replicates for each condition; $\chi 2$ test for no response or swimming response. (C) The data presents the reaction time of each larva in response to the touch stimulus across three groups: the vehicle group (n=56 larvae), the $(AUUUU)_7$ group (n=53 larvae), and the $(AUUUC)_{58}$ group (n=50 larvae), with a total of six experimental replicates for each condition; Kruskal-Wallis test. (D) The data outlines the swimming duration of each animal following the touch stimulus across three groups: the vehicle group (n=29 larvae), the $(AUUUU)_7$ group (n=31 larvae), and the $(AUUUC)_{58}$ group (n=29 larvae), all derived from six experimental replicates; Kruskal-Wallis test. Data are shown as mean ± standard deviation.

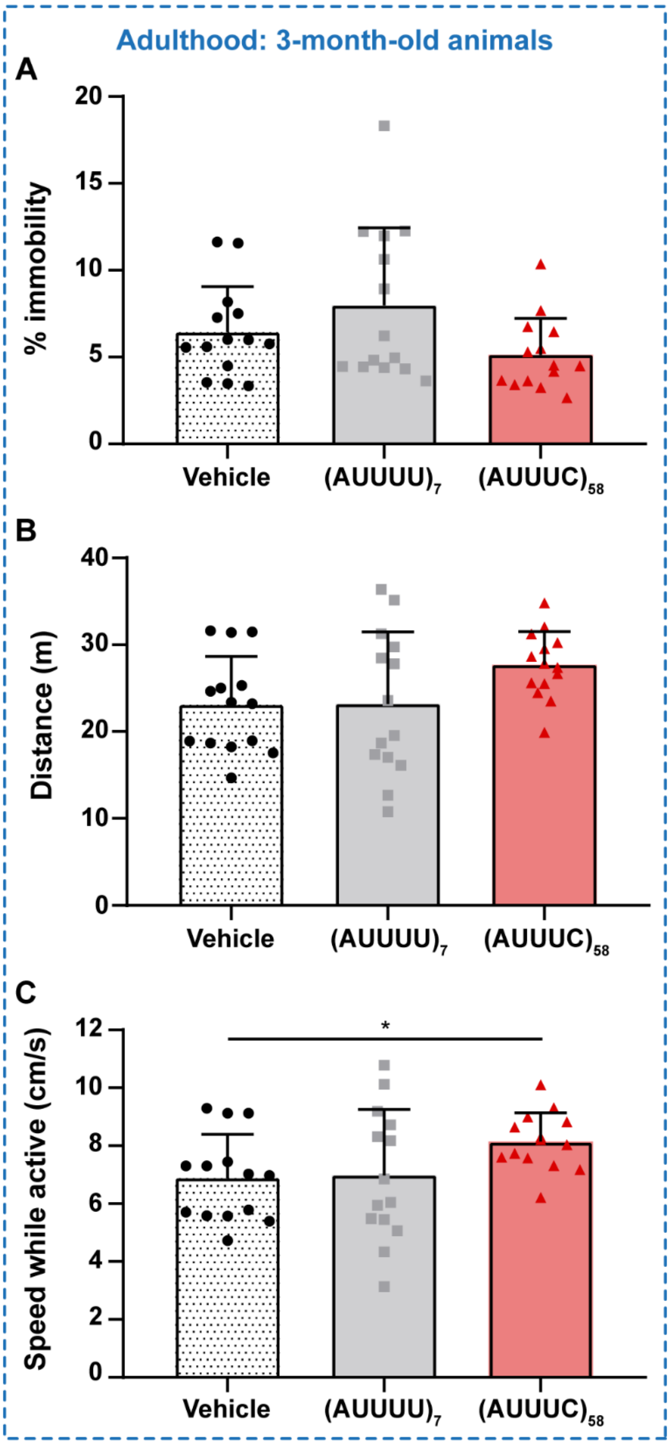


**Fig. S6 The 3-month-old (AUUUC)$_{58}$ zebrafish exhibited no motor impairment.** The novel diving tank test was conducted on microinjected animals with vehicle, (AUUUU)$_7$, or (AUUUC)$_{58}$ at 3 months of age, with a sample size of 14 animals per condition. (A) Analysis of the percentage of immobility (Kruskal-Wallis test). (B) Analysis of the total swimming distance (One-way ANOVA). (C) Measurement of the average swimming speed (*$p<0.05$, One-way ANOVA following Welch correction and Dunnett's T3 post-hoc test). Data are shown as mean ± standard deviation.

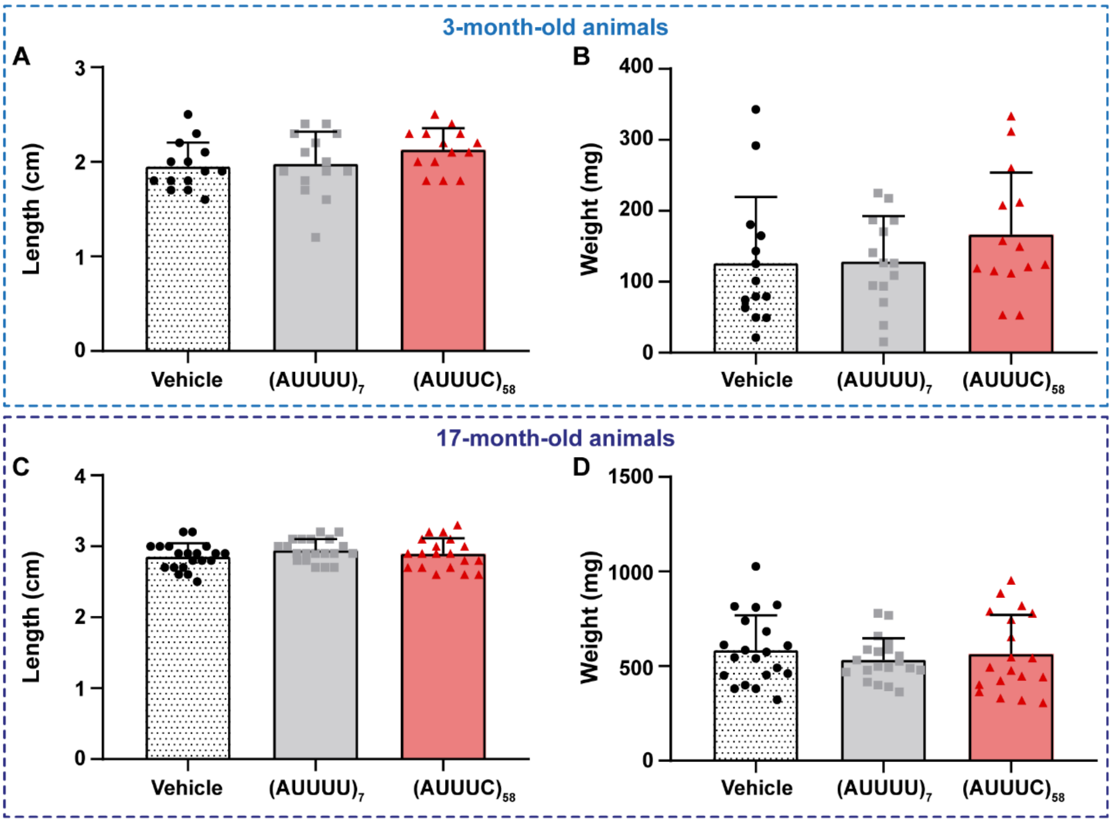


**Fig. S7 Comparison of body size and weight in 3- and 17-month-old animals indicates no significant differences.** The following samples size were used: n=14 animals per condition at 3 months, and n=20 animals for the vehicle and $(AUUUU)_7$ conditions, and n=19 animals for the $(AUUUC)_{58}$ condition at 17 months. The top section presents data for 3-month-old animals, while the bottom section features data for 17-month-old animals. Zebrafish length at 3 months (A) and 17 months (C) was analyzed using a One-way ANOVA, which yielded no significant results. The weight of zebrafish assessed at 3 months (B) and 17 months (D) was evaluated using the Kruskal-Wallis test and One-way ANOVA test, respectively, showing no significant differences. Data are represented as mean ± standard deviation.

**Table S1** Maximum Z-scores for RBPs predicted to bind SCA37 $(AUUUC)_{58}$ RNA, as determined by the RBPmap webserver.

| RBP | Non-pathogenic RNA | SCA37 RNA |
|---|---|---|
| A1CF | 3.267 | 3.267 |
| ANKHD1 | 2.551 | 2.551 |
| BOLL | 3.829 | 3.829 |
| BRUNOL4 | 3.865 | 3.865 |
| BRUNOL5 | 3.85 | 3.85 |
| BRUNOL6 | 3.16 | 3.16 |
| CELF1 | 3.629 | 3.629 |
| CNOT4 | 2.436 | 2.436 |
| CPEB1 | 3.29 | 3.29 |
| CPEB2 | 3.929 | 3.929 |
| CPEB4 | 3.696 | 3.696 |
| DAZ3 | 3.721 | 3.721 |
| DAZAP1 | 2.903 | 2.903 |
| EIF4G2 | 2.727 | 2.727 |
| ELAVL4 | 3.67 | 3.67 |
| EWSR1 | 1.991 | 1.991 |
| FUBP1 | 3.44 | 3.44 |
| FUBP3 | 3.562 | 3.562 |
| FXR1 | 1.7 | 1.7 |
| G3BP2 | 2.053 | 2.053 |
| HNRNPA0 | 3.18 | 3.18 |
| HNRNPA1 | 3.311 | 3.311 |
| HNRNPA1L2 | 1.887 | 1.887 |
| HNRNPA2B1 | 2.745 | 2.745 |
| HNRNPC | 3.634 | 3.634 |
| HNRNPCL1 | 3.658 | 3.658 |
| HNRNPD | 2.988 | 2.988 |
| HNRNPDL | 3.152 | 3.152 |
| HNRNPF | 2.361 | 2.361 |
| HNRNPH1 | 1.93 | 1.93 |
| HNRNPL | 3.013 | 3.013 |
| HNRNPM | 2.159 | 2.159 |
| HNRNPU | 3.031 | 3.031 |
| HNRPLL | 2.861 | 2.861 |
| HuR | 3.857 | 3.857 |
| IGF2BP1 | 1.761 | 1.761 |
| IGF2BP2 | 3.466 | 3.466 |
| IGF2BP3 | 3.373 | 3.373 |
| KHDRBS1 | 2.928 | 2.928 |
| KHDRBS2 | 2.788 | 2.788 |

**Table S1** (continued)

| RBP | Non-pathogenic RNA | SCA37 RNA |
|---|---|---|
| KHDRBS3 | 2.972 | 2.972 |
| KHSRP | 3.282 | 3.282 |
| LIN28A | 2.5 | 2.5 |
| MATR3 | 2.703 | 2.703 |
| MBNL1 | 2.903 | 2.903 |
| MSI1 | 2.476 | 2.476 |
| NOVA1 | 2 | 4 |
| NUPL2 | 2.636 | 2.636 |
| PABPC1 | 3.235 | 3.235 |
| PABPC3 | 3.216 | 3.216 |
| PABPC4 | 3.327 | 3.327 |
| PABPC5 | 3.068 | 3.068 |
| PABPN1 | 2.528 | 2.528 |
| PABPN1L | 3.094 | 3.094 |
| PCBP1 | 2.044 | 2.044 |
| PCBP2 | 3.037 | 3.037 |
| PCBP3 | 3.635 | 3.635 |
| PCBP4 | 2.806 | 2.806 |
| PRR3 | 2.274 | 2.274 |
| PTB3 | 3.296 | 3.296 |
| PTBP3 | 3.337 | 3.337 |
| PUF60 | 3.403 | 3.403 |
| PUM1 | 3.211 | 3.211 |
| PUM2 | 2.966 | 2.966 |
| QKI | 2.207 | 2.207 |
| RALY | 4.17 | 4.17 |
| RBFOX1 | 3.039 | 3.039 |
| RBM15B | 4.319 | 4.319 |
| RBM23 | 2.9 | 2.9 |
| RBM24 | 4.323 | 4.323 |
| RBM28 | 2.514 | 2.514 |
| RBM38 | 3.974 | 3.974 |
| RBM41 | 3.42 | 3.42 |
| RBM42 | 2.811 | 2.811 |
| RBM45 | 3.089 | 3.089 |
| RBM47 | 2.78 | 2.78 |
| RBM6 | 3.139 | 3.139 |
| RBMS1 | 3.836 | 3.836 |
| RBMS2 | 3.19 | 3.19 |
| RBMS3 | 3.656 | 3.656 |
| RC3H1 | 3.535 | 3.535 |
| SAMD4A | 1.716 | 1.716 |

**Table S1** (continued)

| RBP | Non-pathogenic RNA | SCA37 RNA |
|---|---|---|
| SART3 | 3.235 | 3.235 |
| SF1 | 2.72 | 2.72 |
| SFPQ | 2.731 | 2.731 |
| SNRNP70 | 1.767 | 1.767 |
| SNRPA | 2.237 | 2.237 |
| SRSF1 | 2.839 | 2.839 |
| SRSF10 | 2.787 | 2.787 |
| SRSF11 | 1.868 | 1.868 |
| SRSF2 | 3.375 | 3.375 |
| SRSF4 | 1.842 | 1.842 |
| SRSF5 | 2.763 | 2.763 |
| SRSF8 | 3.463 | 3.463 |
| SRSF9 | 2.743 | 2.743 |
| TARDBP | 2.99 | 2.99 |
| TIA1 | 4.127 | 4.127 |
| TRA2A | 2 | 2 |
| TRNAU1AP | 3.988 | 3.988 |
| TUT1 | 1.836 | 1.836 |
| U2AF2 | 3.571 | 3.571 |
| UNK | 2.672 | 2.672 |
| YBX1 | 3.571 | 3.571 |
| YBX2 | 3.608 | 3.608 |
| ZC3H14 | 4.062 | 4.062 |
| ZCRB1 | 2.531 | 2.531 |
| ZFP36 | 4 | 4 |
| ZNF326 | 2.975 | 2.975 |
| ZNF638 | 3.108 | 3.108 |

RBP – RNA-binding protein; NOVA1 highlighted in red; $p<0.05$

### Supplementary Movies

**Movie 1.** Representative video file capturing spontaneous tail coil movements in a zebrafish embryo microinjected with vehicle control.

**Movie 2.** Representative video file capturing spontaneous tail coil movements in a zebrafish embryo microinjected with the non-pathogenic $(AUUUU)_7$ RNA control.

**Movie 3.** Representative video file capturing spontaneous tail coil movements in a zebrafish embryo microinjected with the pathogenic $(AUUUC)_{58}$ RNA.
